# Regulatory Plasticity and Metabolic Trade-offs Drive Adaptive Evolution of Alternative Flagellar Configurations in *Pseudomonas aeruginosa*

**DOI:** 10.1101/2025.07.29.667523

**Authors:** Anali Migueles Lozano, Merrill Asp, Sofia T. Rocha, Jiaqi Li, Georgia Fanouraki, Aden D. Sun, Lichun Zhang, Jacob R. Waldbauer, Jiarong Hong, Abhishek Shrivastava, Jing Yan, Sampriti Mukherjee

**Affiliations:** Department of Molecular Genetics & Cell Biology, University of Chicago, Chicago, IL, USA; Department of Microbiology, University of Chicago, Chicago, IL, USA; Duchossois Family Institute, University of Chicago, Chicago, IL, USA; Department of Molecular, Cellular and Developmental Biology, Yale University, New Haven, CT, USA; Yale Quantitative Biology Institute, Yale University, New Haven, CT, USA; School of Life Sciences, Arizona State University, Tempe, AZ, USA; Biodesign Institute, Arizona State University, Tempe, AZ, USA; Department of Mechanical Engineering, University of Minnesota, 111 Church Street SE, Minneapolis, MN, USA; Department of the Geophysical Sciences, University of Chicago, Chicago, IL, USA

## Abstract

Evolutionary constraints governing flagellar number in bacterial pathogens remain poorly understood. While related *Pseudomonas* species are hyperflagellated, *P. aeruginosa* maintains strict monoflagellation through the FleQ-FleN regulatory circuit. Here, we demonstrate that FleN dosage is essential for maintaining monoflagellation and bacterial fitness. Wild-type *P. aeruginosa* consistently displayed unipolar monoflagellation, while Δ*fleN* mutants developed over two-to-five flagella per cell in uni- or bipolar arrangements. Hyperflagellated Δ*fleN* cells exhibited severe fitness defects including reduced growth rates, attenuated virulence in nematode infection models, and competitive disadvantages in co-culture experiments. Remarkably, Δ*fleN* cells rapidly evolved suppressor mutations in *fleQ* that partially restored growth and motility without always restoring monoflagellation. Five independent suppressor alleles mapped to critical FleQ functional domains (four in the AAA+ ATPase domain, one in the DNA-binding domain), suggesting reduced protein activity that rebalances the disrupted regulatory circuit. Single-cell motility analysis revealed that suppressor strains exhibit heterogeneous swimming dynamics, with subpopulations achieving wild-type speeds despite carrying multiple flagella. Proteomic analysis demonstrated that hyperflagellation triggers extensive cellular reprogramming beyond flagellar components, affecting metabolic pathways, stress responses, and signaling networks. While hyperflagellated cells suffered complete loss of pathogenicity in animal infection models, environmental selection under viscous conditions could drive wild-type cells to evolve enhanced motility through specific *fleN* mutations. These findings suggest that bacterial flagellar regulatory circuits function as evolutionary capacitors, normally constraining phenotypic variation but enabling rapid adaptation to alternative motility configurations when environmental pressures exceed the performance limits of standard monotrichous flagellation.

**SIGNIFICANCE STATEMENT:** Bacterial flagella are extracellular appendages that rotate to propel the cell and enable swimming motility. While some bacteria have multiple flagella, many pathogenic species like *Pseudomonas aeruginosa* have just one. Surprisingly, mutants of *P. aeruginosa* with multiple flagella performed worse, i.e., they grew more slowly, were less infectious in laboratory animals, and were outcompeted by wild-type bacteria. Even when some mutant bacteria evolved compensatory changes, they still struggled compared to single-flagellum bacteria. This reveals an important evolutionary trade-off: while multiple flagella might seem advantageous for movement, having just one flagellum allows bacteria to grow faster and cause more severe infections. This plasticity likely explains why *P. aeruginosa* is so successful both in the environment and as a human pathogen.

## INTRODUCTION

The specification of appendage number and location is a fundamental biological requirement across all domains of life. Through evolutionary processes, organisms develop specific arrangements and functions (such as feeding and locomotion) of these appendages to optimize survival in their particular environments (Panganiban et al., 1997; Shubin et al., 1997). Evolutionary changes in appendage number are common, even within the same phylogenetic order, for instance, only 40% of Phasmatodea insects (stick insects) possess full wings, with the remainder being partially or completely wingless (Ogden & Whiting, 2003; Zeng et al., 2020, 2023). This precise maintenance of appendage patterns across generations proves equally vital for microorganisms such as bacteria, particularly for their various motility mechanisms: swimming, swarming, gliding, or twitching. The locomotion and maneuverability of microorganisms are central to their fitness, with species-specific motility differences determining population and community-level outcomes as movement capabilities fundamentally shape how cells navigate their environments in search of resources and favorable conditions (Lambert et al., 2006, 2011; Nolan et al., 2018; Toutain et al., 2005). Flagellar motility, in particular, enables rapid swimming and precise chemotactic responses, allowing bacteria to detect and move toward nutrients or away from toxins along chemical gradients (Hintsche et al., 2017; Köhler et al., 2000; Matilla et al., 2022; Paul et al., 2010). This ability to respond to environmental cues through directed movement influences microbial distribution patterns, biofilm formation, and interspecies competition. Beyond nutrient acquisition, flagellar motility plays critical roles in host colonization, pathogenesis, and dispersal, directly impacting transmission dynamics and infection processes (Gode-Potratz et al., 2011; Jain & Kazmierczak, 2014; Ozer et al., 2021; van Ditmarsch et al., 2013). However, there are significant energetic costs of construction and operation of flagella due to the elaborated building of this multi module nanomachine (Bhattacharyya et al., 2024; Ferreira et al., 2019; Lisevich et al., 2025; Schavemaker & Lynch, 2022).

A bacterial flagellum is assembled from over 30 proteins in three architectural domains: the basal body anchoring the flagellum to the cell envelope, the hook universal joint, and the helical propeller-like filament (Chevance & Hughes, 2008; Chilcott & Hughes, 2000; Mondino et al., 2022; Rossmann & Beeby, 2018). Bacteria exhibit diverse flagellar arrangements including monotrichous (single polar), lophotrichous (multiple at one pole), peritrichous (distributed around cell), and amphitrichous (at both poles), that produce distinct swimming patterns (run-tumble, run-reverse, wrapping), diversifying exploration strategies and resource access (Grognot & Taute, 2021; Hintsche et al., 2017; Kinosita et al., 2020; Vater et al., 2014). The peritrichous arrangement is exemplified by *E. coli*, which typically possesses 6-10 flagella randomly distributed across its cell surface, enabling the characteristic run-and-tumble motility pattern essential for chemotaxis and environmental exploration. In the polar flagellates, diversity in flagellar number is exemplified within Pseudomonas: *P. aeruginosa* maintains a single unsheathed polar flagellum, *P. putida* typically possesses 5-7 polar flagella per cell, *P. fluorescens* exhibits strain-specific variations from 2 flagella (MFE01) to single flagellum (SBW25) to 7 flagella (C7R12), while *P. syringae* averages 2.7 flagella per cell with notable stochastic variation and bimodal expression patterns (Nogales et al., 2015). However, key questions remain: why do bacterial species exhibit such flagellar variation, what evolutionary and ecological factors drive monotrichous versus lophotrichous systems, and how do bacteria balance flagellar biosynthesis and operation costs against mobility advantages in different environments?

Here, we addressed these questions using the monotrichous versatile bacterium *Pseudomonas aeruginosa* that can be either pathogenic in humans (de Sousa et al., 2021; Jurado-Martín et al., 2021) or beneficial/harmful in plants (Chahtane et al., 2018; Ghazi Faisal, 2024), with flagellar motility being crucial for its colonization capabilities. Flagellar motility in *P. aeruginosa* directly correlates with increased pneumonia and burn wound sepsis mortality rates (Arora et al., 2005; Hatano et al., 1996; Landsperger et al., 1994; Luzar et al., 1985). Unlike related Pseudomonas species with multiple flagella, monotrichous *P. aeruginosa* is thought to navigate using a “run-reverse” strategy (Vater et al., 2014). Flagellar assembly is regulated through a sophisticated four-tiered hierarchical cascade centered on the transcriptional regulator FleQ: FleQ (class I) activates class II genes in a σ54-dependent manner; FleR (class II) activates class III genes for hook-basal body assembly; once this structure is complete, FlgM is exported, releasing FliA (σ28) to activate class IV genes encoding the flagellar filament and motor proteins (Bouteiller et al., 2021; Nandini Dasgupta et al., 2003). FleQ activity is precisely controlled by the anti-activator FleN (Fig. 1A) (Chanchal et al., 2021; Dasgupta & Ramphal, 2001; Dasgupta et al., 2000; Torres-Sánchez et al., 2023), which binds to FleQ’s ATPase domain, and by cyclic di-GMP (c-di-GMP), which serves as a master switch between motility and biofilm formation (Baraquet & Harwood, 2013); (Hickman & Harwood, 2008; Jiménez-Fernández et al., 2016). In this study, we investigated the fitness consequences of hyperflagellation, i.e., whether there is an evolutionary trade-off between flagellar number and growth/virulence. We demonstrate that although *P. aeruginosa* can develop multiple flagella, this hyperflagellation comes with significant fitness costs, including reduced growth, competitiveness, and virulence. This suggests that *P. aeruginosa* has evolved to maintain regulatory plasticity in its flagellar assembly cascade as an adaptive strategy for its particular ecological niches and pathogenic lifestyle.

**Fig. 1:**
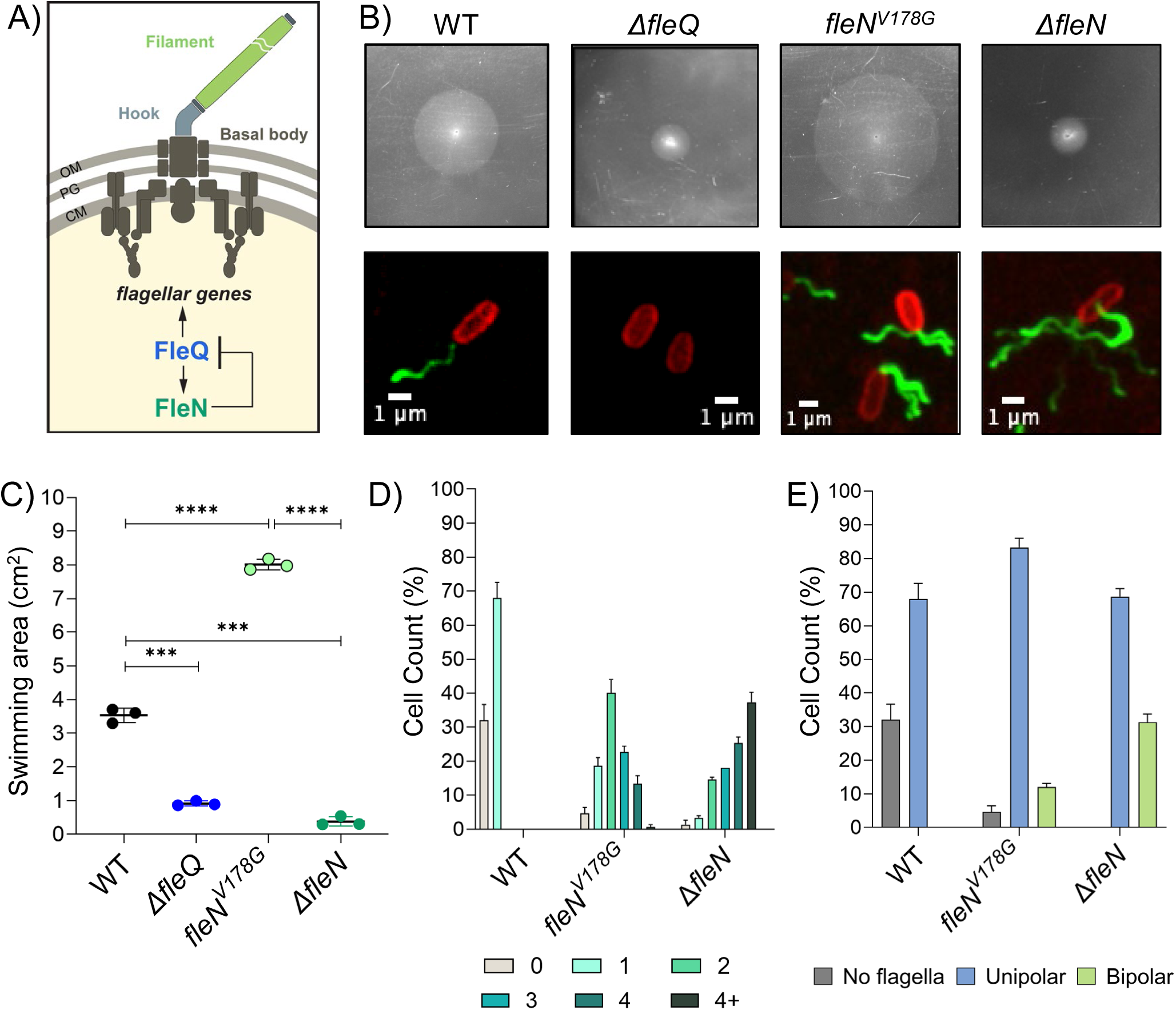
Flagellar number and patterning in WT and *fleN* mutant strains. A) Schematic of the conserved flagellar architectural domains and FleQ-FleN regulatory axis. FleQ is the master transcriptional factor of flagellar genes, and its activity is directly inhibited by the antiactivator FleN. FleQ promotes the expression of FleN. B) Top row: Representative images of swimming phenotype in low agar concentration plates (0.3%) of (from left to right) WT, Δ*fleQ*, *fleN^V178G^* and Δ*fleN* strains. Bottom row: Representative images of flagellar patterning of (from left to right) WT, Δ*fleQ*, *fleN^V178G^* and Δ*fleN*. Flagellar filament and membrane stained using Alexa488 (green) and membrane dye (FM4-64), respectively. C) Swimming area measurement of WT, Δ*fleQ*, *fleN^V178G^* and Δ*fleN* strains. Error bars represent the SD of three biological replicates. Statistical significance was determined using t-test pairwise comparisons in GraphPad Prism software. **** P<0.0001, *** P<0.001 D) Percentage of cells exhibiting varying number of flagella during exponential growth phase (OD_600_ 0.5) in WT, Δ*fleQ*, *fleN^V178G^*and Δ*fleN* strains; 50 cells per replicate, n=3. Grey: non-flagellated, Cyan: 1, Turquoise: 2, Dark turquoise: 3, Emerald green: 4, Dark green: More than 4 flagella per cell. E) Percentage of cells with different flagellar placement from the cells in panel D).

## RESULTS

### Wild-type *P. aeruginosa* PA14 exhibits robust monotrichous flagellation

Our first goal was to systematically define flagellar number and arrangement in wild-type (WT) *P. aeruginosa* UCBPP-PA14 (hereafter called PA14). To visualize and count the number of completed flagella, we introduced into the *fliC* gene, encoding for the filament protein flagellin, a cysteine mutation (*fliC^T394C^*) that allowed labeling with thiol-reactive fluorescent Alexa Fluor 488 dye without affecting swimming motility (Fig.1B). We monitored filament number using confocal microscopy at different growth stages (Fig. 1B,D, E, Supplemental fig. 1). During early exponential phase (OD_600_ 0.25), cells displayed mixed flagellation patterns: 51.33% monotrichous, 48% non-flagellated, and rarely (0.66%) multiple flagella clustered at one pole while from exponential through stationary phase (OD_600_ 0.5-1.5), monotrichous flagellation predominated (68%-82%). We conclude that WT PA14 exhibits robust monotrichous flagellation pattern under planktonic culture conditions in the laboratory.

*P. aeruginosa* encounter dramatically different physical environments throughout its complex lifecycle—from the watery surfaces in natural habitats to the viscous mucus of cystic fibrosis airways (Ambreetha & Balachandar, 2022; Chahtane et al., 2018; Crone et al., 2020; Hatano et al., 1996; Jurado-Martín et al., 2021; Moustafa et al., 2024). This environmental heterogeneity creates fluctuating demands on bacterial motility systems, where optimal flagellar configurations must balance swimming efficiency against metabolic costs under varying physical constraints (Hook et al., 2019; Kojima et al., 2020; Lisevich et al., 2025; Mahenthiralingam et al., 1994; Molina et al., 2020; Moustafa et al., 2024; Wu et al., 2024). To investigate whether environmental pressures could drive evolution of alternative flagellar configurations, we subjected WT *P. aeruginosa* PA14 to selection on viscous swimming medium (0.35% agar compared to standard 0.3%) that increases motility demands. After extended incubation (48 hours at 37°C), rare but consistent outgrowth flares emerged from the primary inoculation zones, indicating cells with improved swimming ability under viscous conditions. Sequencing analysis of independent isolates revealed a striking pattern: all mutants harbored the identical mutation, *fleN^V178G^*. Characterization of PA14 *fleN*^V178G^ cells confirmed enhanced swimming motility compared to WT while assessment of flagellar number and polarity showed increased flagellar numbers per cell and a subpopulation of cells exhibiting amphitrichous hyperflagellation (Fig. 1B-E). This remarkable convergence on a single amino acid, i.e., position 178 in FleN represents a critical regulatory hotspot enabling evolutionary fine-tuning of flagellar output.

The discovery of the naturally evolved FleN^V178G^ variant prompted us to investigate the broader evolutionary potential of the FleN-FleQ regulatory circuit. To explore whether dramatic perturbations might reveal the full range of accessible phenotypes and the mechanistic basis of FleN function in constraining alternative flagellar configurations, we constructed an in-frame marker less deletion of *fleN* and quantified the resulting phenotypes. In the absence of FleN, FleQ activity is unchecked, giving rise to cells with hyperflagellation phenotype (Fig1B, E and Supplemental figure 1). At the early growth stage (OD_600_ 0.25), there are cells having zero (1.33%), one (10.6%), two (16.66%), three (22.66%), four (21.33%) and more than four (27.33%) flagella (Supplemental fig. 1). From exponential growth through stationary phase (OD_600_ 0.5-1.5), the fraction of cells having more than four flagella became the predominant phenotype (37.33-50.66%). Furthermore, *ΔfleN* cells displayed three different types of flagella arrangements: monotrichous (single polar flagellum), amphitrichous (flagella at both poles), and lophotrichous (cluster of flagella at one end), each at different frequencies, lophotrichous being the predominant one (<50% in every growth stage) (Fig. 1B-E). Through exponential growth, the fraction of cells having flagella at both poles increased; from 16.66-31.33% during the exponential phase (OD_600_ 0.25 and 0.5), to 42.66% at the stationary phase (OD_600_ 1.5).

Despite producing multiple flagella, Δ*fleN* cells showed severely compromised swimming motility in soft agar assays, with swimming zone diameters decreased by approximately 70% compared to WT (Fig. 1B,C). This counterintuitive result demonstrated that flagellar number alone does not determine swimming performance – instead, proper flagellar arrangement and coordination are essential for efficient propulsion. Furthermore, this finding provided crucial context for understanding why the V178G mutation, which increases flagellar number only modestly, provides adaptive advantages while absence of FleN impairs function. To confirm that the observed phenotypes resulted specifically from *fleN* loss, we complemented Δ*fleN* cells using an arabinose-inducible *fleN* construct integrated at the chromosomal *attB* site. Arabinose titration experiments (0.006-0.5%) revealed a complex dose-response relationship with distinct threshold effects (Fig. 2A). Low arabinose concentrations (0.01%) restored swimming motility to WT levels, demonstrating functional complementation (Fig 2A). However, flagellar number and patterning remained partially altered compared to WT cells, with 18.4% bearing more than four flagella compared to <5% in WT cells (Fig. 2B, C, D). Flagellar polarity between the arabinose induced (unipolar 82.97%, bipolar 12.91%) and uninduced (unipolar 84.66%, bipolar 15.33%) conditions had a subtle decrease on bipolar flagella. Notably, high arabinose concentrations (0.5%) completely abolished swimming motility and triggered a dramatic shift toward non-flagellated cells (62% with zero flagella, 30.7% with one flagellum) (Fig. 2B, C, D). This bifurcation demonstrates that FleN overexpression blocks flagellar assembly entirely, establishing that precise FleN dosage is critical for normal flagellar regulation. The narrow concentration window supporting normal motility indicates that the FleN-FleQ regulatory circuit operates near a critical threshold, making it sensitive to regulatory perturbations.

**Fig. 2:**
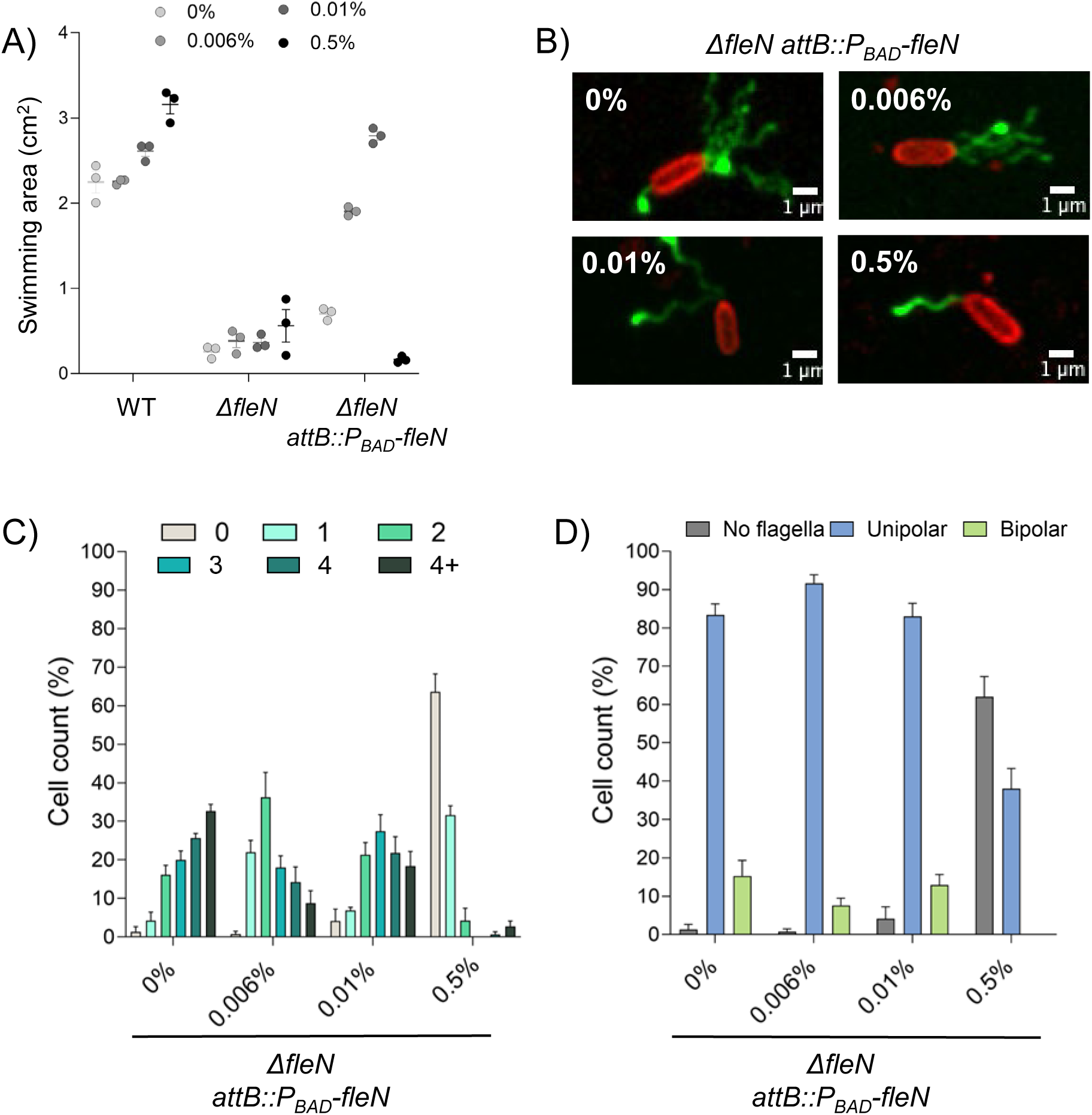
FleN dosage controls flagellar number and swimming motility. A) Swimming area measurement of WT, Δ*fleN* and *ΔfleN attB::P_BAD_fleN* strains, using 0%, 0.006%, 0.01% and 0.5% concentrations of arabinose. Error bars represent the SD of three biological replicates. B) Representative images of flagellar patterning of *ΔfleN attB::P_BAD_fleN* strain at indicated arabinose concentrations. Flagellar filament and membrane stained using Alexa488 (green) and membrane dye (FM4-64), respectively. C) Percentage of cells exhibiting varying number of flagella during exponential growth phase (OD_600_ 0.5) in *ΔfleN attB::P_BAD_fleN* strain at the indicated arabinose concentrations, 50 cells per replicate, n=3. Grey: non-flagellated, Cyan: 1, Turquoise: 2, Dark turquoise: 3, Emerald green: 4, Dark green: More than 4 flagella per cell. D) Percentage of cells with different flagellar placement from the cells in panel C).

### Selection pressure drives convergent evolution of *fleQ* suppressor mutations

To investigate how Δ*fleN* cells might naturally overcome their motility defects, we performed adaptive evolution experiments by plating Δ*fleN* cultures on 0.3% swimming agar and selecting for enhanced motility over 48 hours at 37°C. This selection consistently yielded distinct outgrowth flares containing cells with improved swimming ability compared to the parental Δ*fleN* strain, suggesting rapid adaptive evolution. We isolated and characterized 10 independent suppressor strains that demonstrated reproducibly enhanced swimming motility (Supplemental table 1). Sequencing analysis revealed a striking pattern: all these suppressors (hereafter referred to as ***s***uppressor ***o***f ***f***leN, i.e., *son*) contained single nucleotide changes exclusively in *fleQ*, suggesting that the FleN-FleQ regulatory circuit represents an evolutionary hotspot for motility adaptation (Fig. 3A). Four mutations (L327M (son1), R340H (son2), N367S (son3), V383G (son4)) clustered within the AAA+ ATPase domain essential for ATP hydrolysis and protein-protein interactions, while one mutation (E442Q, son5) localized to the C-terminal helix-turn-helix DNA-binding domain required for target gene recognition (Fig. 3A). Based on this distribution, we hypothesized that suppression of motility defect occurred through impaired FleQ function. To test whether these mutations caused partial loss of FleQ function, we introduced each *fleQ* allele into WT PA14 and assessed swimming motility. All five mutations either decreased motility compared to WT or showed no significant effect, confirming partial loss-of-function properties (Supplemental Fig. 2). However, when *fleN* was deleted in each *fleQ^son^* mutant background, motility defects were enhanced relative to WT, demonstrating that these mutations provide suppression specifically in the sensitized Δ*fleN* context (Fig. 3B, Supplemental Fig. 2). Western blot analysis using custom raised antibodies against FleQ showed that the protein levels of the WT and FleQ^son^ mutants were comparable, thereby ruling out protein stability defects (Supplemental Fig. 4). We conclude that these suppressor mutations restore regulatory balance by reducing FleQ activity to compensate for the absence of FleN-mediated inhibition.

**Fig. 3:**
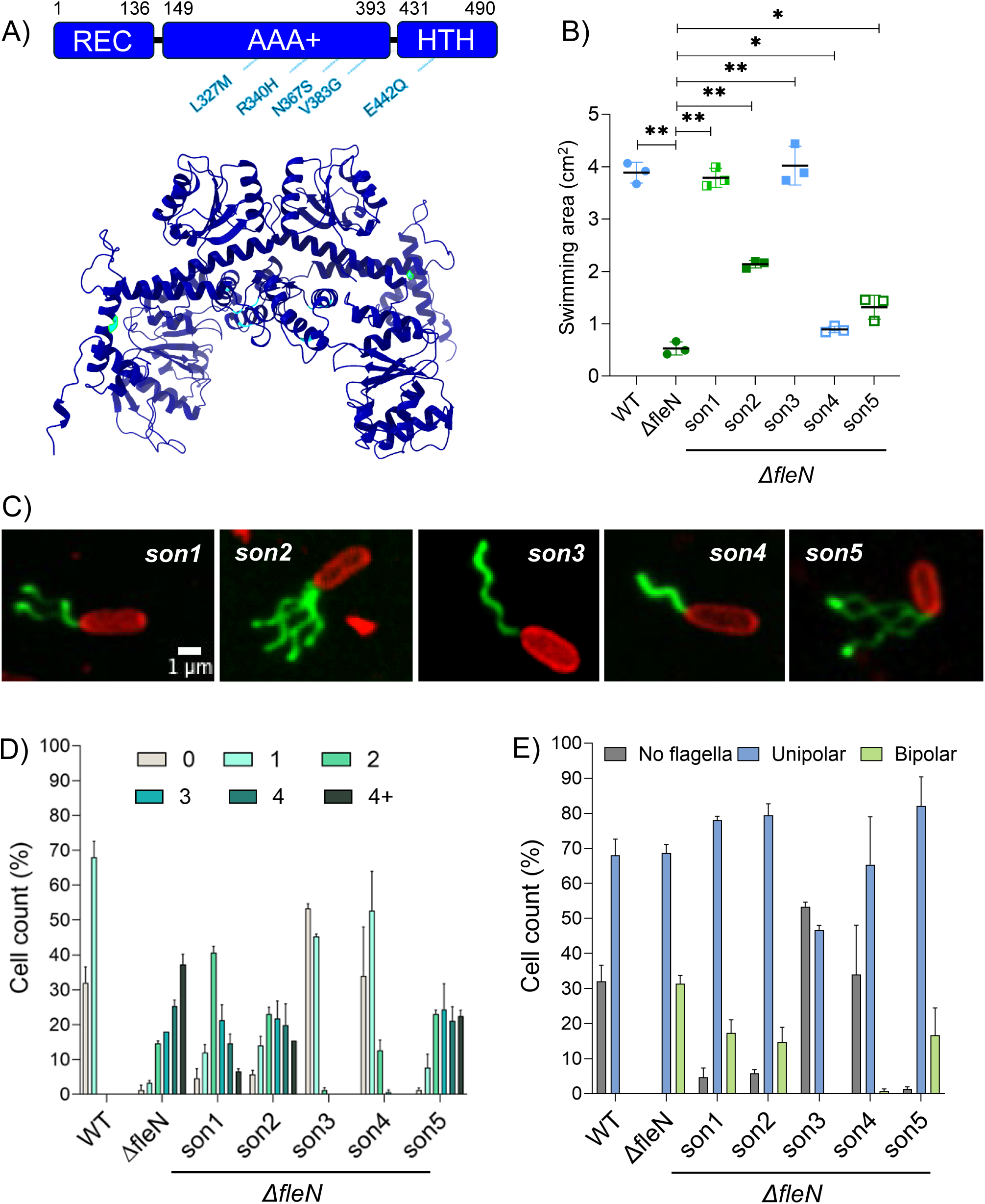
Δ*f*leN cells develop suppressor mutations in *fleQ* that restore motility but not number of flagella. A) Box (up) and Alphafold3 (down) representation of FleQ protein. In the Box diagram and Alphafold3 models, Δ*fleN* suppressor mutations in *fleQ* are denoted in cyan. Alphafold3 model of FleQ dimer (navy blue). B) Swimming area measurement of the indicated strains. Error bars represent the SD of three biological replicates. Statistical significance was determined using t-test pairwise comparisons in GraphPad Prism software. ** P <0.005, * P<0.01. C) Representative images of flagellar patterning of the indicated strains: *son1* (*fleQ^L327M^*), *son2* (*fleQ^R340H^*), *son3* (*fleQ^N367S^*), *son4* (*fleQ^V383G^*), *son5* (*fleQ^E442Q^*). Flagellar filament and membrane stained using Alexa488 (green) and membrane dye (FM4-64), respectively. D) Percentage of cells exhibiting varying number of flagella during exponential growth phase (OD_600_ 0.5) in the indicated strains, 50 cells per replicate, n=3. Grey: non-flagellated, Cyan: 1, Turquoise: 2, Dark turquoise: 3, Emerald green: 4, Dark green: More than 4 flagella per cell. E) Percentage of cells with different flagellar placement from the cells in panel D).

Quantitative flagellar number and arrangement analysis of the five suppressor strains during exponential growth revealed two phenotypic classes of suppressors – class-1 mutants exemplified by *son3* and *son4*, showed near-complete restoration of WT flagellar numbers with predominantly monotrichous cells, while class-2 mutants exemplified by son1, son2 and son5 retained varying degrees of hyperflagellation (Fig. 3C, D, E, Supplemental Fig. 3). Nonetheless, all suppressor strains shared common features distinguishing them from the ancestral Δ*fleN* strain: reduced frequency of bipolar flagellation and decreased proportion of cells with more than four flagella (Fig. 3E). This convergent phenotypic pattern indicates that excessive flagellar assembly, particularly at multiple cellular poles, represents the primary constraint limiting motility in hyperflagellated cells. We note that motility in 0.3% agar plates showed a disconnection with flagellar number; for instance, *son4* cells despite achieving WT flagellar counts, demonstrated minimal improvement in swimming motility compared to the parental Δ*fleN* strain. Conversely, *son1* cells maintained significant hyperflagellation yet achieved swimming speeds comparable to WT.

### Single-cell analysis reveals population heterogeneity in swimming dynamics

To understand how different flagellar configurations affect individual cell motility, we tracked the centroid positions of individual cells swimming in liquid media across PA14, Δ*fleN*, and *son2* strains. Over one thousand quantitative, three-dimensional trajectories were captured with digital in-line holography (Sheng et al., 2006), with several hundred per strain, as detailed in Methods. This revealed typical instantaneous speeds of swimming cells and the shape of their trajectories while tracked over a 15 second time frame. The effective speed (displacement divided by time within the tracked window) reflects both the instantaneous speed and how straight or meandering the cell’s path is. WT PA14 cells exhibited characteristic straight swimming patterns with typical mean speeds of 10 μm/s and the top 5% of mean speeds in excess of 24 μm/s, under the particular conditions we tested. Across all strains, a consistently observable subpopulation remained less motile (Fig. 4A). The Δ*fleN* mutant strain showed dramatically altered swimming dynamics, with highly helical and meandering trajectories that reduced typical mean speeds to about 6 μm/s, with the top 5% of mean speeds only 15 μm/s or above (Fig. 4B). Analysis of trajectory shape across the population revealed significant heterogeneity, with a subset of Δ*fleN* cells achieving WT swimming patterns (relatively straight trajectories at 10 μm/s or more) despite the overall population deficit – consistent with the observation that Δ*fleN* cells show reduced but not abolished motility on swimming plates. The helical and meandering trajectories were almost never observed in the WT population, suggesting that miscoordination between multiple flagella may cause less efficient swimming patterns. This finding indicates that hyperflagellation generates phenotypic variation within isogenic populations, generating cells with distinct motility capabilities.

**Fig. 4:**
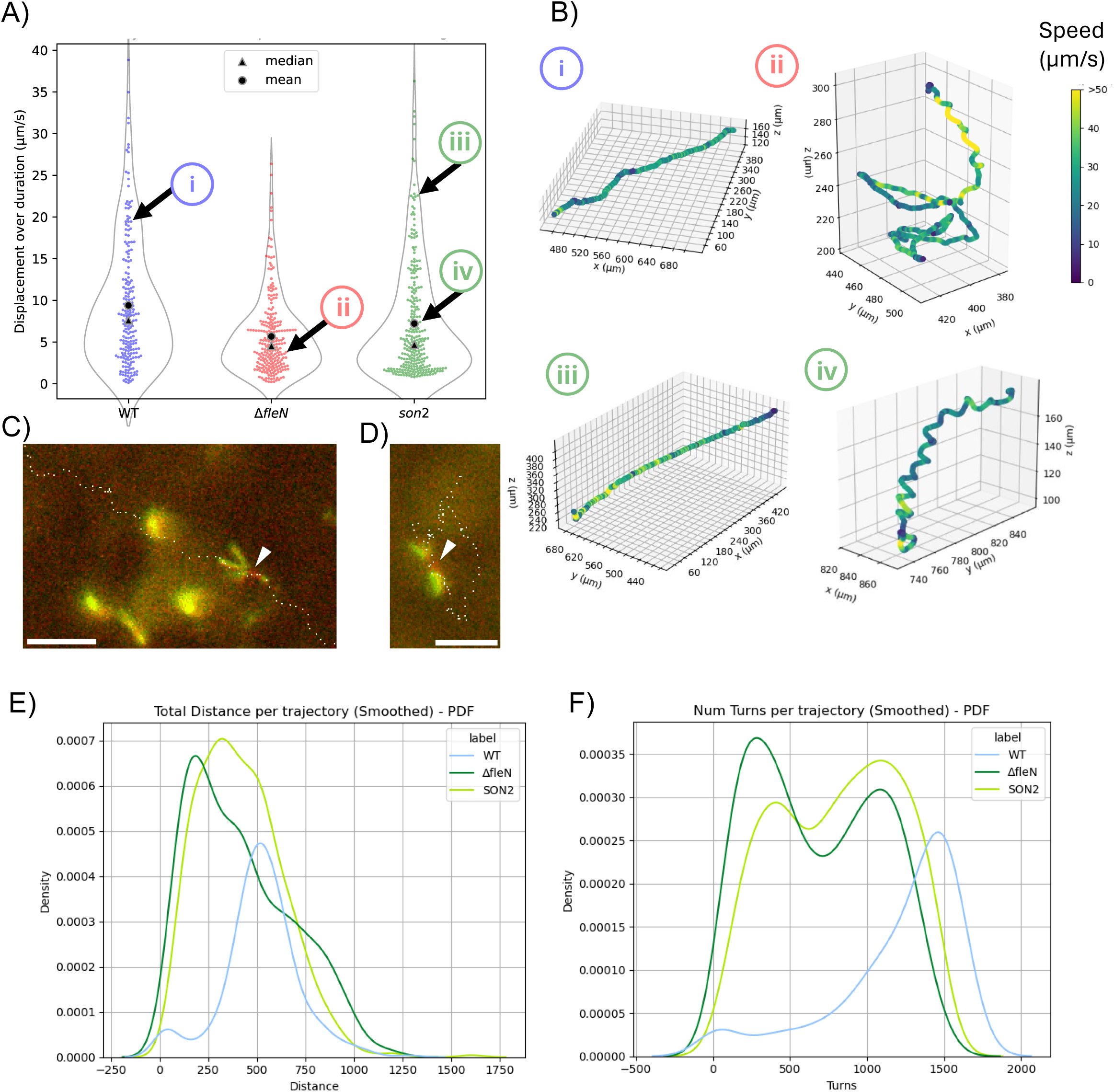
3D Holographic tracking of single cell dynamics shows heterogenous swimming behaviors. A) Mean speed (displacement over time) is calculated for individual cell trajectories captured swimming for up to 15 seconds. Each point represents an individual cell, with data collected across two biological replicates. B) Three-dimensional trajectories are displayed for four individual cells with mean speeds pointed out as i-iv in panel A. Color indicates instantaneous speed. i) A straight WT trajectory, showing typical high mean speed behavior. ii) A low mean speed Δ*fleN* trajectory, showing both a helical path (upper) and a meandering path (lower). iii) A high mean speed *son2* trajectory, showing recovery of swimming in a straight line. iv) A low mean speed *son2* trajectory showing strong helicity. C) A *son2* cell (cell body in red, constitutively fluorescent) with unbundled flagella (green) on its left side, with both channels imaged simultaneously while the cell is swimming. A small point of green on the right side indicates flagella on the opposite pole, possibly wrapped around the cell. White dots show the progress of the cell body over 3.3 seconds of swimming, displaying a directed helical trajectory. Scale bar 10 µm. D) Another *son2* cell with flagella active on both poles, undergoing a meandering trajectory, as shown by the white dots that mark the cell body position during 3.8 seconds of swimming. Scale bar 10 µm. E-F) Distribution of the single cells’ behaviors across the different strains in terms of E) Distance per trajectory and F) Number of turns per trajectory. Smoothed using kernel density estimation (KDE).

We further supplemented the centroid trajectory data with direct observation of swimming cell’s flagella by employing simultaneous dual-color fluorescence microscopy and capturing videos, with details included in Methods. This produced several illustrative examples of how hyperflagellation impacts individual cell motility, including confirming that helical trajectories are consistent with hyperflagellated cells (Fig. 4C) and the most meandering trajectories typically belong to bipolar hyperflagellated cells (Fig. 4D). Flagella dynamically bundle and unbundle during swimming, affecting direction and speed, although tight flagellar bundles are observed to not be necessary for swimming as in other multiflagellated species such *E. coli*. Hyperflagellated cells were occasionally observed swimming faster than typical WT cells, briefly achieving speeds up to 50 μm/s in one instance. Cells with multiple flagella swim best when only flagella on one pole are active, although cells with flagella on the opposite pole were observed wrapping around the cell to achieve persistent motion as well. Overall, increasing the number of flagella drastically increases the variety of swimming modes available to cells, which can switch stochastically over time between different behaviors even for individual cells. The *son2* suppressor mutant demonstrated its own distinct distribution of behaviors, with helical trajectories and typical mean speeds similar to Δ*fleN* but expanded subpopulations of both fast-swimming cells and slow-swimming cells, pushing the top 5% of mean speeds back up to 22 μm/s or more. Notably, analysis of the total distance per trajectory and number of turns per trajectory for WT, Δ*fleN and son2* mutants showed that the shift toward WT swimming behavior in the *son2* mutant (Fig. 4E, F). Taken together, we conclude that rather than uniform improvement across the population, the *son2* mutant appears to have increased the fraction of cells capable of overcoming hyperflagellation constraints.

### Hyperflagellation imposes significant metabolic costs

To investigate the broader physiological consequences of hyperflagellation, we performed comparative growth analysis and discovered that Δ*fleN* cells exhibited significantly slower growth kinetics compared to WT (Fig. 5A). While WT cultures reached stationary phase (OD₆₀₀ = 1.5) within 20 hours, Δ*fleN* cultures achieved only OD₆₀₀ = 1.0 in the same timeframe (Figure 5A). Serial dilution experiments confirmed equal viability between strains, but Δ*fleN* colonies remained smaller after extended incubation, indicating sustained growth defects (Supplemental Fig. 5A). Furthermore, cell morphological analysis during exponential phase revealed that Δ*fleN* cells displayed heterogeneous sizes, with subpopulations of cells either larger or smaller than WT (Supplemental Fig. 5B). This size heterogeneity correlates with the observed growth rate reduction and suggests that hyperflagellation affects fundamental cellular processes beyond flagellar assembly and motility. Since FliF is required early in flagellar assembly, *fliF* deletion blocks production of all downstream flagellar components, substantially reducing the total metabolic investment in flagellar biosynthesis. To test whether growth defects resulted from the metabolic burden of flagellar overproduction, we constructed deletion strains removing *fliF* (encoding a basal body component, 64.0 kDa) from WT and Δ*fleN* backgrounds. Absence of FliF significantly improved growth compared to Δ*fleN* alone (p < 0.0001), though not to WT levels (Fig. 5C). We note that the hyperswimmer and hyperflagellated *fleNV^178G^* mutant shows no significant growth difference when compared to WT (Fig. 5B). Nonetheless, the growth dynamics of suppressors of *ΔfleN* correlate with number of flagella per cell: single flagellated suppressors (son3 and son4) recover growth dynamics similar to WT while hyperflagellated suppressors (son1, son2, son5) exhibit growth dynamics that range in between WT and *ΔfleN* gowth curves (Fig. 5D). The partial growth restoration confirms that flagellar overproduction contributes significantly to the fitness costs observed in Δ*fleN* cells, though additional factors beyond simple metabolic burden are likely involved.

**Fig. 5:**
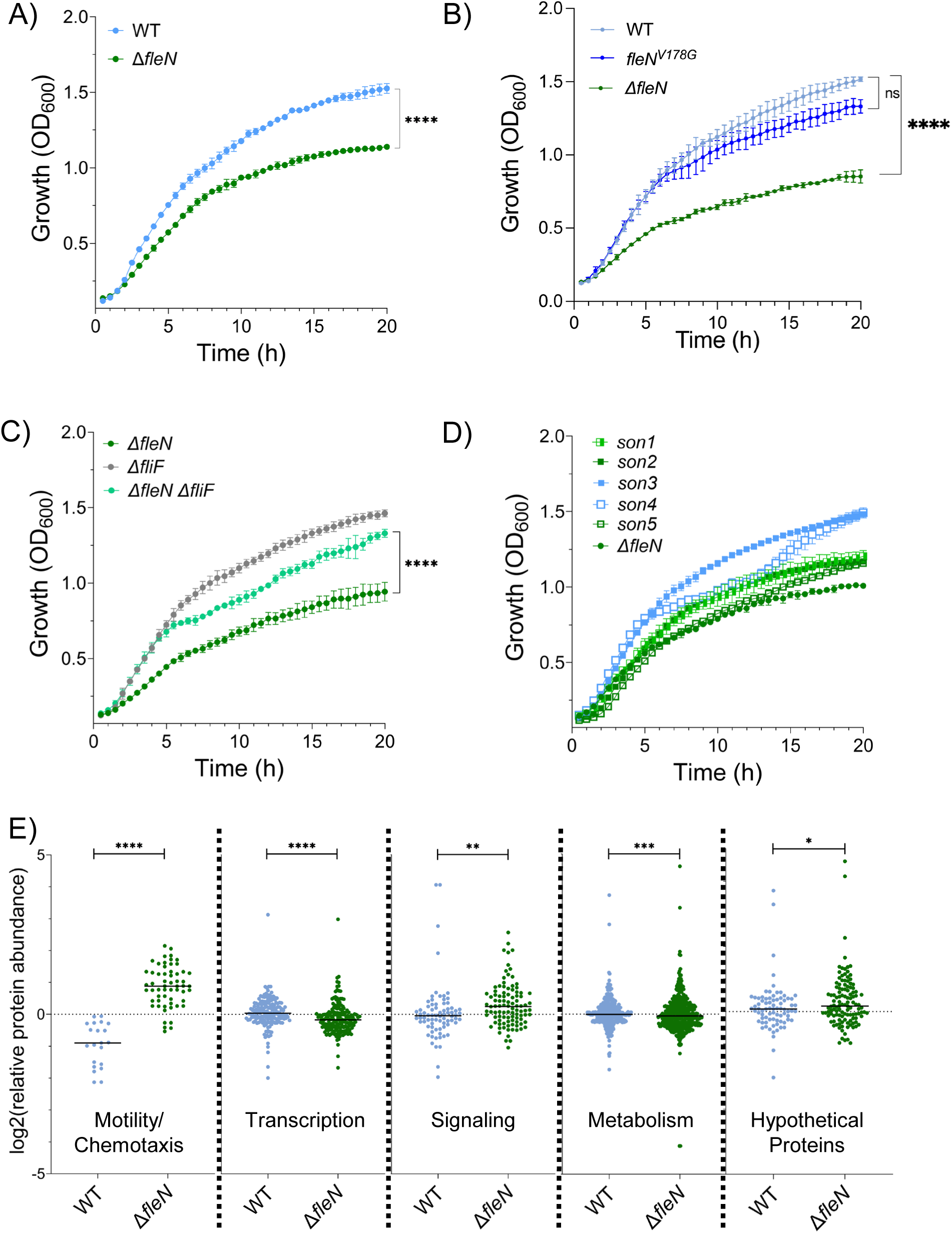
Δ*f*leN mutant exhibits a growth defect compared to WT. A) Growth curve of WT and Δ*fleN* cells, n=3. B) Growth curve of WT, *ΔfleN* and *fleN^V178G^* strains, n=3. C) Growth curve of *ΔfliF*, *ΔfleN* and *ΔfleNΔfliF* strains, n=3. D) Growth curve of Δ*fleN* suppressors, *son1-5*. For panels A-C, statistical test: nonparametric Mann Whitney test, significant p-values are less than 0.01. E) Proteomics analysis of WT and Δ*fleN* strains. t-test Non-parametric, Kolmogorov-Smirnov (KS) test. Significant p-values are less than 0.01.

To comprehensively assess the molecular consequences of hyperflagellation, we performed quantitative proteomics using the diDO-IPTL method (Waldbauer et al., 2017) to compare protein abundances between WT and Δ*fleN* cells during exponential growth (Figure 5E). We classified differentially abundant proteins into functional categories based on Gene Ontology terms: Motility and Chemotaxis, Metabolism and Biosynthesis, Signaling, Transcription, Cell Division and DNA Replication, Stress Response and Protection, and Unclassified proteins. As expected, proteins in the Motility and Chemotaxis category showed significant overabundance in Δ*fleN* cells, including core flagellar structural components (FlgE, FliF, FliC, FliM) and motor proteins (MotC), confirming the hyperflagellation phenotype at the molecular level (Figure 5E).

However, extensive changes occurred across all other functional categories, revealing that FleN deletion triggered broad cellular reprogramming beyond flagellar assembly. Several metabolic enzymes increased significantly, including cysteine synthase A (PA14_29110, upregulated ∼eight folds), aconitate hydratase (PA14_44290, upregulated ∼thirteen folds) and acyl carrier protein AcpP (PA14_25670, upregulated >3 folds), while cytochrome c4 (PA14_72460) showed over four-fold reduction. Signaling pathway proteins demonstrated coordinated upregulation, particularly phenazine biosynthesis enzymes (PhzE, PhzB, PhzC, PhzF) and two-component regulatory systems (CheY, FleR). Furthermore, proteins such as bacterioferritin (PA14_18670, upregulated 4.8 folds) and catalase (PA14_09150, upregulated ∼four folds) were more abundant in the Δ*fleN* mutant compared to WT, suggesting activation of systems linked to stress responses. Transcriptional regulatory proteins displayed particularly striking alterations: while transcription was downregulated overall in the Δ*fleN* mutant compared to WT, among the most abundant proteins in Δ*fleN* cells were PA14_48770 (putative OsaR ortholog involved in osmoregulation), transcription termination factor Rho (PA14_69190), the stationary phase sigma factor RpoS (PA14_17480), and the flagellar sigma factor FliA (PA14_45630). This pattern suggests that hyperflagellation activates alternative transcriptional programs, possibly as stress responses to the altered cellular state. In addition, several hypothetical proteins including PA14_03350, PA14_44080, PA14_50570, PA14_58330, PA14_69090 were greater than five folds more abundant in the Δ*fleN* mutant compared to WT. Taken together, these global proteomic changes reflect altered membrane composition requirements for supporting multiple flagellar assemblies or metabolic adaptations to the energetic costs of hyperflagellation.

### Hyperflagellation severely reduces competitive fitness and pathogenic capacity

To assess the ecological consequences of hyperflagellation, we performed direct competition experiments between WT and Δ*fleN* cells in liquid culture. Starting with equal cell densities, WT cells achieved complete dominance within 17 hours, reaching >10:1 ratios in colony-forming units (Fig. 6A, Supplemental Fig. 5C). This competitive disadvantage confirms that the growth defects observed in monoculture translate to severe fitness costs under competitive conditions. Furthermore, investigation of the consequences of hyperflagellation on type IV pili mediated twitching motility showed inverse correlation such that the Δ*fleN* mutant was severely impaired in twitching (Fig. 6B). Consistent with this, analysis of the five *fleQ* suppressor strains revealed that suppressors that restored monotrichous flagellation (*son3* and *son4*) maintained WT level twitching, while those retaining hyperflagellation (*son1, son2,* and *son5*) showed severely reduced twitching despite their improved swimming ability (Fig. 6B). We conclude that hyperflagellation hinders twitching motility.

**Fig. 6:**
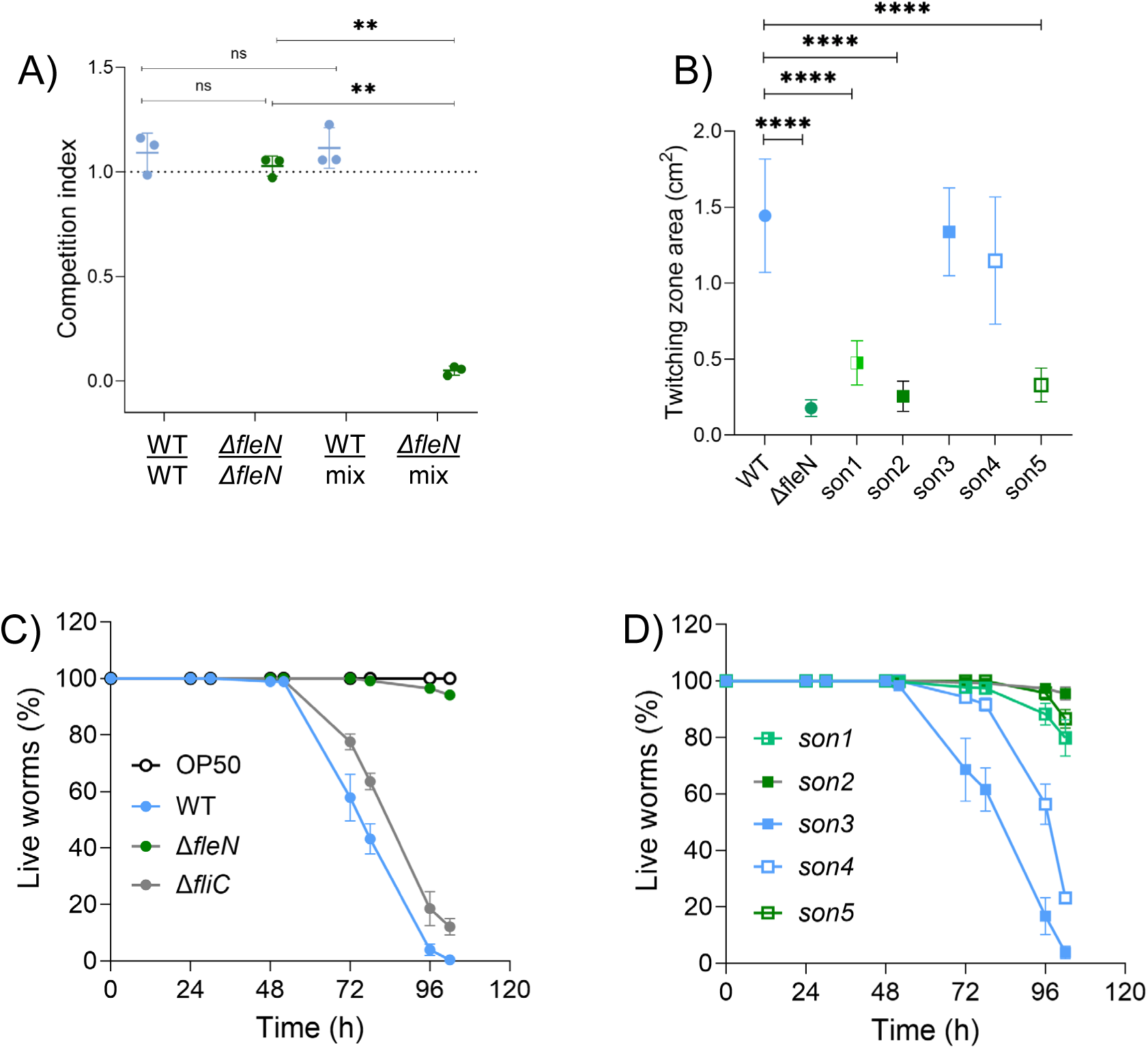
Hyperflagellated strains exhibit competitive disadvantage and decreased twitching and virulence. A) Competition between WT and *ΔfleN* cells in shaken cultures. Competition index (CI) is the ration between number of cells of each strain at the end of the assay relative to number of cells of each strain at the beginning of the assay, n=3. CI∼1= No difference, CI<1=Mutant cells decrease, CI>1=Mutant cells increase. Statistical tests: WT/*ΔfleN and WTmix/ΔfleNmix* unpair t-test. WT/WTmix and *ΔfleN/ΔfleNmix* parametric Ratio paired t-test. B) Twitching motility of WT, *ΔfleN and ΔfleN* suppressor strains. Error bars represent the SD of three biological replicates. Statistical significance was determined using t-test pairwise comparisons in GraphPad Prism software. **** P<0.0001 C-D) Worm slow-killing assay where *C. elegans* were applied to lawns of *E. coli* OP50 and the indicated WT and mutant *P. aeruginosa* strains. Error bars represent SEM of three independent experiments.

Next, we investigated pathogenic capacity using the *Caenorhabditis elegans* slow-killing assay, a well-established model for *P. aeruginosa* virulence (Tan et al., 1999; Vasquez-Rifo et al., 2019; Xue et al., 2024). WT PA14 demonstrated full pathogenic potential, killing 100% of exposed worms within 96 hours (Fig. 6C). In striking contrast, the Δ*fleN* mutant showed virtually no pathogenicity, with worm survival rates comparable to non-pathogenic *E. coli* OP50 control bacteria (Fig. 6C). To determine whether this virulence defect resulted from impaired motility or other consequences of FleN deletion, we tested a non-motile Δ*fliC* mutant lacking the flagellar filament entirely. Remarkably, non-flagellated cells retained complete pathogenic capacity, killing worms with kinetics identical to WT (Fig. 6C). This result demonstrates that flagellar motility or the flagellum itself is not required for PA14 pathogenesis in this model system, implying that hyperflagellation specifically, rather than loss of motility/flagella per se, impairs virulence mechanisms. Consistent with this, analysis of the five *fleQ* suppressor strains revealed a clear correlation between flagellar assembly phenotype and pathogenic capacity. Suppressors that restored monotrichous flagellation (*son3* and *son4*) maintained full virulence, while those retaining hyperflagellation (*son1, son2,* and *son5*) remained avirulent despite their improved swimming ability (Fig. 6D). We conclude that hyperflagellation and its associated metabolic costs, rather than the loss of motility, impairs pathogenic processes.

## DISCUSSION

Our systematic investigation of flagellar regulatory circuit disruption reveals that *P. aeruginosa* maintains unexpected evolutionary flexibility in flagellar assembly, enabling rapid adaptation to alternative motility configurations through targeted regulatory mutations. Our systematic characterization of flagellar patterns in WT PA14 confirms its predominant monotrichous nature, with cells displaying either a single polar flagellum or no flagellum, particularly during active growth. This binary flagellation pattern supports previous observations of *P. aeruginosa* as a polar monoflagellated bacterium (Amako & Umeda, 1982; Dasgupta et al., 2003). However, our work demonstrates that this pattern is not simply a default state but rather the result of active regulation through the FleN-FleQ system, a regulatory axis that represents a tunable system for flagellar control. Our complementation experiments with titratable FleN expression revealed a remarkable “cliff-edge” relationship between FleN levels and swimming motility. The observation that intermediate FleN levels (0.01% arabinose) restored optimal swimming despite maintaining altered flagellation patterns suggests that swimming efficiency depends on precise regulation of the FleN-FleQ axis rather than strictly adhering to monotrichous flagellation, as further confirmed by the heterogeneous swimming behaviors in 3D single-cell trajectories.

The suppressor mutations that naturally emerged in the Δ*fleN* background provide further evidence for the critical role of the FleN-FleQ regulatory axis. These mutations, primarily localized to the AAA+ and HTH domains of FleQ, represent a compensatory mechanism that partially restores the regulatory balance disrupted by FleN deletion. The spectrum of flagellar phenotypes observed among these suppressors - ranging from near-WT monotrichous patterns to intermediate hyperflagellation - demonstrates the remarkable plasticity of the flagellar regulatory system. Intriguingly, the dissociation between flagellar pattern and swimming efficiency observed in some suppressor strains (e.g., *son4* showing WT-like flagellation but minimal swimming improvement, while *son1* maintains hyperflagellation yet swims efficiently and *fleN^V178G^* mutant is hyperflagellated and hyperswimmer) suggests that additional factors beyond mere flagellar number and arrangement influence motility. These might include coordinated flagellar rotation, chemotaxis system integration, or metabolic efficiency.

On the technical side, we have introduced the use of digital in-line holography (DIH) to visualize the trajectories of *P. aeruginosa* cells in three dimensions. Compared to other commonly used particle tracking techniques, DIH has the advantages of straightforward implementation and suitability to track cells precisely even in a high-density culture (Yu et al., 2014). Using DIH, we observed three broad categories of trajectory for individual *P. aeruginosa* cells: straight (most common in WT), meandering (observed with bipolar hyperflagellated cells), and helical (observed in Δ*fleN* and suppressor mutant populations). The relative sizes of the subpopulations expressing these motility patterns are affected by the FleN-FleQ regulatory axis, creating opportunities for the most motile subpopulations to be selected for by environmental conditions. The increased diversity in swimming behaviors for hyperflagellated strains may lead to the evolutionary potential for the hyperflagellated phenotype to be selected, as is naturally seen in several other Pseudomonas species.

Our findings demonstrate that hyperflagellation in *P. aeruginosa* PA14 imposes substantial fitness costs that extend beyond impaired motility. The reduced growth rate, altered cell morphology, and compromised pathogenicity of the Δ*fleN* strain highlight the multifaceted consequences of dysregulated flagellar production. The restoration of growth in the Δ*fleN*Δ*fliF* strain suggests that the fitness burden stems primarily from overproduction of basal body components rather than filament proteins, consistent with the hierarchical nature of flagellar gene expression (Dasgupta et al., 2003; Kutsukake et al., 1990). The fitness costs associated with hyperflagellation provide an explanation for the evolutionary selection of monotrichous flagellation in *P. aeruginosa*, particularly when the multiple flagella do not coordinate to increase swimming thrust as we observed in the Δ*fleN* mutant. In natural environments where resources are limited and competition is intense, the energetic burden of producing and maintaining multiple flagella would place hyperflagellated variants at a significant disadvantage. Indeed, our competition experiments demonstrate that WT PA14 rapidly outcompetes the Δ*fleN* strain in co-culture, supporting the notion that resource efficiency is a critical factor in the evolution of flagellar patterns. The differential pathogenicity observed between monotrichous and hyperflagellated PA14 strains in the *C. elegans* infection model adds another dimension to the fitness equation. The finding that flagellar motility is not essential for pathogenesis aligns with previous reports on clinical isolates (Amiel et al., 2010; Wolfgang et al., 2004), but the striking loss of virulence in hyperflagellated strains suggests that flagellar dysregulation impacts pathogenicity through mechanisms beyond motility. These might include altered immune recognition, disrupted biofilm formation, or compromised secretion of virulence factors, aspects that warrant further investigation.

Despite the predominance of monotrichous flagellation in *P. aeruginosa*, our discovery of the *fleN^V178G^* mutation that confers hyperflagellation, enhanced swimming in increased viscosity with no significant growth defect suggests that alternative flagellar arrangements might be advantageous under specific environmental conditions. We note that this mutation has been reported previously (Deforet et al., 2014; van Ditmarsch et al., 2013), but our study suggests that while unipolar hyperflagellation can confer increased motility we find precipitous decrease in motility in strains with >4 flagella at one pole. This finding resonates with ecological studies showing that bacteria can adapt their motility systems to diverse habitats (Ferreira et al., 2019; Toft & Fares, 2008). Similarly, *P. fluorescens* engineered to lack flagellar motility via deletion of *fleQ* consistently regained swimming ability by evolving the homologous protein NtrC to function as an activator of flagellar gene transcription (Taylor et al., 2015, Shepherd et al., 2023). Furthermore, the rarity of such adaptations in our experimental setup, compared to the frequent emergence of suppressors in the Δ*fleN* background, suggests that transitions from monotrichous to alternative flagellar patterns are tightly controlled and occur only under strong selective pressure. This evolutionary conservatism likely reflects the generally favorable cost-benefit ratio of monotrichous flagellation across the diverse environments that *P. aeruginosa* typically inhabits. Synteny analysis (Supplemental Fig. 6) shows high conservation of flagellar gene clusters between monotrichous (*P. aeruginosa*) and lophotrichous (*P. putida, P. syringae*) Pseudomonas. Despite *P. aeruginosa* being traditionally associated as a human colonizer, it can also succeed in environments commonly colonized by species such as *P. putida* and *P. syringae*, in the phyllosphere and rhizosphere (Ambreetha & Balachandar, 2022; Chahtane et al., 2018; Ghazi Faisal, 2024; Liu et al., 2018; Molina et al., 2020; Sánchez-Gil et al., 2023; K. Yu et al., 2019). It is worth mentioning that within the rhizosphere, there have been reports of Pseudomonas communities expressing different levels of flagellar genes (López-Pagán et al., 2025), concordant with the results found in our study about heterogeneity of flagella patterning within a population.

Additionally, sequence alignments of FleQ, FleN and FliC proteins from different Pseudomonads showed high conservation between FleQ and FleN, but not within FliC sequence (Supplemental Fig. 7, 8, 9). These differences in amino acid sequence could be related with evolutionary optimization of bundling, stabilization and coordination of flagellin filaments in lophotrichous Pseudomonas, tailored to the different viscosity levels within the rhizosphere. Taken together, our findings contribute to the broader understanding of bacterial motility by demonstrating that flagellar arrangements represent not merely fixed taxonomic characteristics but adaptable traits under complex regulation, and that regulatory circuits controlling bacterial flagella operate as evolutionary capacitors, generally restricting trait variation but permitting quick adaptive transitions to new movement patterns when environmental stresses exceed what standard monotrichous flagellation can handle. The balance between motility performance, resource efficiency, and pathogenic potential likely varies across ecological niches, potentially explaining the diversity of flagellar patterns observed across bacterial taxa. These findings enhance our understanding of bacterial motility regulation and provide insight into the evolutionary forces shaping flagellar systems across the bacterial kingdom.

## MATERIALS AND METHODS

### Strains and growth conditions

P. aeruginosa UCBPP-PA14 strain was grown in lysogeny broth (LB) (10 g tryptone, 5 g yeast extract, 5 g NaCl per L), in 1% tryptone broth (TB) (10 g tryptone per L), and on LB plates fortified with 1.5% Bacto agar at 37°C. When appropriate, antibiotics were included at the following concentrations: 400 µg/mL carbenicillin, 50 µg/mL gentamycin, 100 µg/mL irgasan, trimethoprim 500 µg/mL. For the CFU quantification, overnight cultures were diluted back to 1:100 into fresh LB and grown over the day until reaching exponential phase (OD_600_ 0.5). Cultures were again diluted back to reach a new concentration of OD_600_ 0.05 and grown for 20hrs. After 20hrs, ten-fold dilutions were plated in LB plates and grown at 37C overnight. Counting of colonies was done manually. For the growth curves, overnight cultures of bacteria were diluted back to OD_600_ 0.05 in 250ul of fresh LB in replicates using a 96well plate. Readings were taken every 30 minutes for 20hrs at 37C with constant shaking. The optical density at 600nm was measured using a Synergy Neo2 microplate reader (Agilent BioTek).

### Strain construction

Strains and plasmids were constructed as described previously (Mukherjee et al., 2017). To construct marker-less in-frame chromosomal deletions in *P. aeruginosa*, DNA fragments flanking the gene of interest were amplified, assembled by the Gibson method, and cloned into pEXG2 (Hmelo et al., 2015). The resulting plasmids were used to transform Escherichia coli SM10λpir and, subsequently, mobilized into *P. aeruginosa* PA14 via biparental mating. Exconjugants were selected on LB containing gentamicin and irgasan, followed by recovery of deletion mutants on LB medium containing 5% sucrose. Candidate mutants were confirmed by PCR. For the construction of *son* mutants, PA14 genomic DNA from the isolated Δ*fleN* suppressors was used as a template. ∼1000bp upstream and downstream of *fleQ* (including *fleQ*) was amplified and cloned into pEXG2. The resulting plasmids were used to transform *E. coli SM10λpir* and, subsequently, mobilized into P. aeruginosa PA14 Δ*fleQ* or Δ*fleN*Δ*fleQ*, respectively.

The complementation strain of *fleN* was cloned into pTJ1-mini-Tn7T-Tp plasmid (Choi, et.al 2006). The resulting plasmids were transformed to Escherichia coli DH5α, and mobilized into P. aeruginosa PA14 via quad-parental mating using as helper strains DH5α/pTNS3 and DH5α/pRK2013 to facilitate Tn7 transposition at the attB site as described here (Choi, et.al., 2006). Insertion of the Tn7 transposon region was confirmed by PCR followed by Sanger sequencing.

### Complementation of FleN

The L(+)-arabinose 99% (AC365180250) was dissolved in miliQ-water and filter sterilized before using it. Cells ablated from *fleN* and carrying a *fleN* insertion under the P_BAD_ were grown in liquid LB overnight without arabinose. For the growth curves, microscopy and swimming assays, arabinose was added at the indicated concentration at the beginning of each experiment.

### Flagella and Membrane Staining

Pseudomonas cells’ flagellin (*fliC*) gene was exchanged with a version of *fliC* (fliCT394C) able to react with Alexa Fluor™ 488 C5 Maleimide from Thermo Fisher (cat. no. A10254). The new strain became the wild-type background used to build all the mutant strains used in this paper. Overnight cultures were diluted 1:100 in 3 mL LB media and then grown over the day at 37°C until reaching the desired OD_600_ (between 0.25 and 1.5). The ends of the pipette tips were cut off consistently for all the staining procedures to minimize flagellar damage from shear. 500 µl of bacterial culture was centrifuged for 1 min at 15,000 rpm, the supernatant was discarded, and the pellet was mixed with Alexa Fluor in 50 µl of 1X PBS buffer. Cells were incubated in the dark for 3 minutes and then 1 mL of 1X PBS was added gently to wash. Cells were then centrifuged for 1 min at 15,000 rpm and the supernatant was discarded. For the membrane staining, similar steps to the flagella staining were used, but using the membrane dye FM464 instead of the Alexa Fluor. Cells were incubated in 50 µl (10 µg/mL) of FM464 (Invitrogen T13320) for 2 min and then washed with 1X PBS. After centrifugation and decanting the supernatant, the pellet was gently resuspended in ∼50 µl of 1X PBS buffer. 10 µl of the sample was placed into a microscopy slide (previously cleaned with 70% isopropanol) and pressed down with a coverslip (previously soaked in poly-L-lysine to immobilize the cells). All microscopy images were taken using the inverted confocal microscope Leica Stellaris 5. Images were then processed using Fiji. Quantification of number and polarity of flagella were done manually, and values were graphed using GraphPad.

### Holographic tracking of cell trajectories in three dimensions

To observe the three-dimensional trajectories of bacteria swimming in liquid media, overnight culture was regrown in LB, cultured at 37°C with shaking for 3.5 hours, until OD_600_ reached approximately 0.8. This was diluted in fresh LB to a concentration with an OD_600_ of 0.002. Samples were prepared by pipetting 20 µL of diluted culture into silicone gaskets (Grace Biolabs) adhered to a cleaned glass coverslip. Digital in-line holography was accomplished with a custom laser illumination setup similar to that described before (Mallery & Hong, 2019) on a Nikon Ti-E using a 4X objective with 1.5X intermediate magnification. Raw holograms were captured for 15 seconds at 120 frames per second on a Teledyne FLIR camera (BFS-U3-32S4M-C) illuminated with a 405 nm laser (Thorlabs) at approximately 5 mW collimated to a beam waist diameter of 5.3 mm. After background subtraction, holograms were processed into X,Y,Z coordinates for the cells using the software RIHVR (Mallery & Hong, 2019)GPU-accelerated with a GeForce RTX 4090. The Python package trackpy (Allan et al., 2024) was used to link the trajectories for further analysis with custom Python code.

### Dual-color flagellar observation assay

To fluorescently label flagella, each cysteine-mutated strain was inoculated from frozen stock into lysogeny broth (LB) and cultured overnight at 30°C with shaking. Overnight culture was regrown in LB, cultured at 37°C with shaking for 3.5 hours, until OD_600_ reached 0.8. Then regrown culture was centrifuged at 4000 rcf for 4 minutes, and the supernatant was removed and replaced with Alexa Fluor C5-maleimide (ThermoFisher) at a concentration of 25 µg/mL diluted In LB. Cells were resuspended by gentle pipetting. This solution incubated at room temperature for 10 minutes before being washed three times by centrifuging cells as before, replacing supernatant with fresh LB, and gently pipette mixing the cells to resuspend. All cell transfers were accomplished with pipette tips that had been pre-cut to widen the opening and prevent flagellar damage due to shear. Cells were transferred into 10 micron-tall PDMS microchannels with glass bottoms for imaging using EPI-illumination on a Nikon Ti2-E. Simultaneous dual color video was achieved with a Cairn OptoSplit II placed at the end of the light path just before the EMCCD, an Andor Ixon life 888.

### Swimming motility assay

The swimming assay was performed as previously published with minor modifications (Ha, D. G., et al., 2014). To the autoclaved solution (0.3% agar media), 200 mL of 5X M8 salts were added, together with 5 mL of 0.2% casamino acids and 40% glucose, respectively, plus 1 mL of MgSO_4_. When appropriate, antibiotics or induction molecules were added at this point. Plates were stacked into piles of 3 and allowed to solidify at room temperature for 4-5 hrs. Overnight cultures (or grown over the day, when specified) were inoculated into the agar by inserting a pipette tip (without reaching the end of the plate) slightly soaked with bacteria. Plates were placed inside containers with soaked paper towels in the base of the container and covering the plates. The plates were placed inside containers with soaked paper towels in the base and top of the container and incubated at 37°C with a water bucket inside of the incubator to maintain humidity. After 21 hrs, plates were imaged using Amersham ImageQuant 800. The swimming area was measured using Fiji and plotted using GraphPad.

### Twitching motility assay

Overnight cultures were inoculated into the flat bottom of a 1.5% agar plate (20 mL each) using skewers. The skewer’s tip was submerged into the overnight cultures and then the tip was pushed through the 20 mL of the solidified agar. Incubation of the plates was done for 48 hrs at 37°C. To stain the twitching area, first a spatula was used to trace along the edge of the plate and lever the agar out of the plate. Next, a thin layer of 0.1% crystal violet was poured on top of the twitching areas (just enough to cover the surface of the plate) and incubated for 20 minutes at room temperature. When ready, the crystal violet was discarded into a waste beaker and then the plate was submerged inside a bowl of clean water, to wash off the excess crystal violet. After letting the plate dry (∼20 minutes), the colorimetric images were collected using the Amersham ImageQuant 800 (Cytiva) imager. Further processing (quantification of twitching area) of the images was done in Fiji with manual tracing of the area. Values were graphed using GraphPad.

### Western Blot Assays

PA14 strains were grown overnight at 37°C in LB medium. The next day, the overnight cultures were diluted back to 1:100 in fresh LB and then grown over the day at 37°C with shaking until reaching OD_600_ 0.5. 1 mL of cell culture was harvested and resuspended in 200 μL of 2X loading buffer (50% 2X Tris buffer (10 mM Tris pH 6.8, 1.5mM bromophenol blue), 45% 2X sample buffer (125 mM Tris pH 6.8, 20% glycerol, 210 mM SDS, 10% BME), 5% BME). The samples were heated for 10 minutes at 95°C. 10 μL samples were loaded onto 12% SDS-PAGE gels and were subjected to electrophoresis.

Proteins were transferred to nitrocellulose membranes overnight (10 V for 16 hrs) at 4°C on a rocking platform. Next day, the membrane was blocked with 5% skim milk in TBS at room temperature for 1 h. Following, an incubation with primary antibodies anti-FleQ and anti-RNAP (Abcam, Inc., 469 Cambridge, UK) at 1:2,500 and 1:10,000 dilution respectively in 5% skim milk in TBS was performed for 2 hrs at room temperature on a rocking platform. Membranes were washed 3 times with TBS-Tween 20 at room temperature for 10 min on a rocking platform. Incubation with the secondary antibody anti-rabbit IgG-Peroxidase produced in goat (Sigma-Aldrich) was followed for 1 h at room temperature. The membranes were washed again 3 times with TBS-Tween 20 for 10 min and subsequently developed with a SuperSignal West Femto Kit (Thermo Scientific) and captured with an Amersham ImageQuant 800 (Cytiva) imager. Quantification of the western blots was performed with Image Lab 6.1 software (Bio-Rad).

### Fluorescent microscopy

Each strain of bacteria was streaked out in LB plates and left at 37°C for 16 hrs, and overnight cultures of single colonies were set. The next day, overnight cultures were diluted back at 1:100 dilution in fresh LB and grown over the day until reaching the desired OD_600_. Unless indicated, 500 μL of the over the day culture was centrifuged for 2 minutes at 15,000 rpm and prepared for flagella staining. 5 μL of the samples were loaded onto a microscope slide and covered with a 1 mm poly-L-Lysine coated coverslip. Micrographs were acquired at 63X magnification with immersion oil in a Stellaris 5 (Leica) microscope and processed using Fiji.

### Nutrient competition

Overnight cultures of bacteria were diluted back (1:100) in fresh LB and then grown over the day until reaching OD_600_ 0.5. To prepare the cells for microscopy, 1 mL of cells of each strain was centrifuged for 2 minutes at 15,000 rpm and the resulting pellet was resuspended in 500 μL of 1X PBS. 5 μL of sample was pipetted onto each microscopy slide. For the competition experiment, in 3 mL of LB, 5 μL of WT and mutant cells (at OD_600_ 0.5) were mixed in a 1:1 ratio. Cultures were incubated while shaking for 18 hrs at 37°C. After the 18 hrs incubation, cultures were diluted back to reach an OD_600_ 0.5 and processed for fluorescence microscopy as previously described. To quantify the cells, 10 areas of interest (30 x 30 µm) per sample, per replica were analyzed using Fiji and manually counted. For the competition index, the number of cells per strain at the end of the experiment (output) were compared to the number of cells of the same strain at the beginning of the experiment (input). Regarding the mixed cultures, the number of cells per strain within the mixed culture at the end of the experiment (output) was compared to the number of cells of the same strain at the beginning of the experiment (input).

### Quantitative Proteomics Analysis

*P. aeruginosa* cell pellets were extracted by heating and vortexing at 95 °C for 20 min in a reducing and denaturing buffer (SDS (1%)/ Tris (200 mM, pH 8.0)/DTT (10 mM)) and cysteine thiols alkylated with 40 mM iodoacetamide. Proteins were then purified by a modified eFASP (enhanced filter-aided sample preparation) protocol (Erde et al. 2014), using Sartorius Vivacon 500 concentrators with a 30 kDa nominal cutoff. Proteins were digested with MS-grade trypsin (37 °C overnight), and peptides were eluted from the concentrator and dried via vacuum centrifugation. Peptides were then isotopically labeled at *N*- and *C*-termini using the diDO-IPTL methodology (Waldbauer et al. 2017) for quantitative proteomic analysis. Briefly, *C*-termini were labeled with oxygen-16 or -18 by an enzymatic exchange in isotopic water of a >98 atom % enrichment, and *N*-termini were labeled with un- or dideuterated formaldehyde via reductive alkylation using sodium cyanoborohydride. Peptide extracts from each sample were split, and aliquots were labeled separately with CD_2_O/H_2_ ^16^O and CH_2_O/H ^18^O; the latter were pooled to serve as a common internal standard for quantification. Aliquots of the ^16^O-labeled peptides and ^18^O-labeled internal standard were mixed 1:1 (v/v) and analyzed by LC−MS for protein expression quantification.

The peptide samples were then separated on a monolithic capillary C18 column (GL Sciences Monocap Ultra, 100 *μ*m I.D. × 200 cm length) with a water−acetonitrile and 0.1% formic acid gradient (2−50% ACN over 180 min) at 360 nL min^−1^ using a Dionex Ultimate 3000 LC system with nanoelectrospray ionization (Proxeon Nanospray Flex source) for LC−MS analysis. Mass spectra were collected on an Orbitrap Elite mass spectrometer (Thermo) operating in data-dependent acquisition mode, with one high-resolution (120,000 *m*/Δ*m*) MS1 parent ion full scan triggering 15 rapid-mode MS2 CID fragment ion scans of selected precursors. Proteomic mass spectral data were analyzed using MorpheusFromAnotherPlace (MFAP) (Waldbauer et al. 2017) with precursor and product ion mass tolerances set to 20 ppm and 0.6 Da, respectively. Static cysteine carbamidomethylation and variable methionine oxidation, *N*-terminal (d4)-dimethylation, and *C*-terminal ^18^O_2_ were included as modifications. The false discovery rate for peptide-spectrum matches was controlled by target-decoy searching to <0.5%. Protein-level relative abundances and standard errors were calculated in *R* using the Arm postprocessing scripts for diDO-IPTL data. (Waldbauer et al. 2017)

## ACKNOWLEDGEMENTS

We thank all members of the Mukherjee lab for thoughtful discussions. Research reported in this publication was supported by the National Science Foundation 2438891&2438892 (to J.Y. and J.H.), the Simons Foundation International (SFI-LS-ECIAMEE-00006634, J.Y.), the National Institute of General Medical Sciences of the National Institutes of Health (NIH) under Grant R35GM150803 to S.M. A.S. and S.T.R. were supported by an NIH award R35GM147131. The content of this study is solely the responsibility of the authors and does not necessarily represent the official views of the funding agencies. The funders had no role in study design, data collection and analysis, decision to publish, or preparation of the manuscript.

## DATA AVAILBILITY

All relevant data are provided within the manuscript and the supplementary information.

## SUPPLEMENTAL FIGURE LEGENDS

**Supplemental Fig.1:**
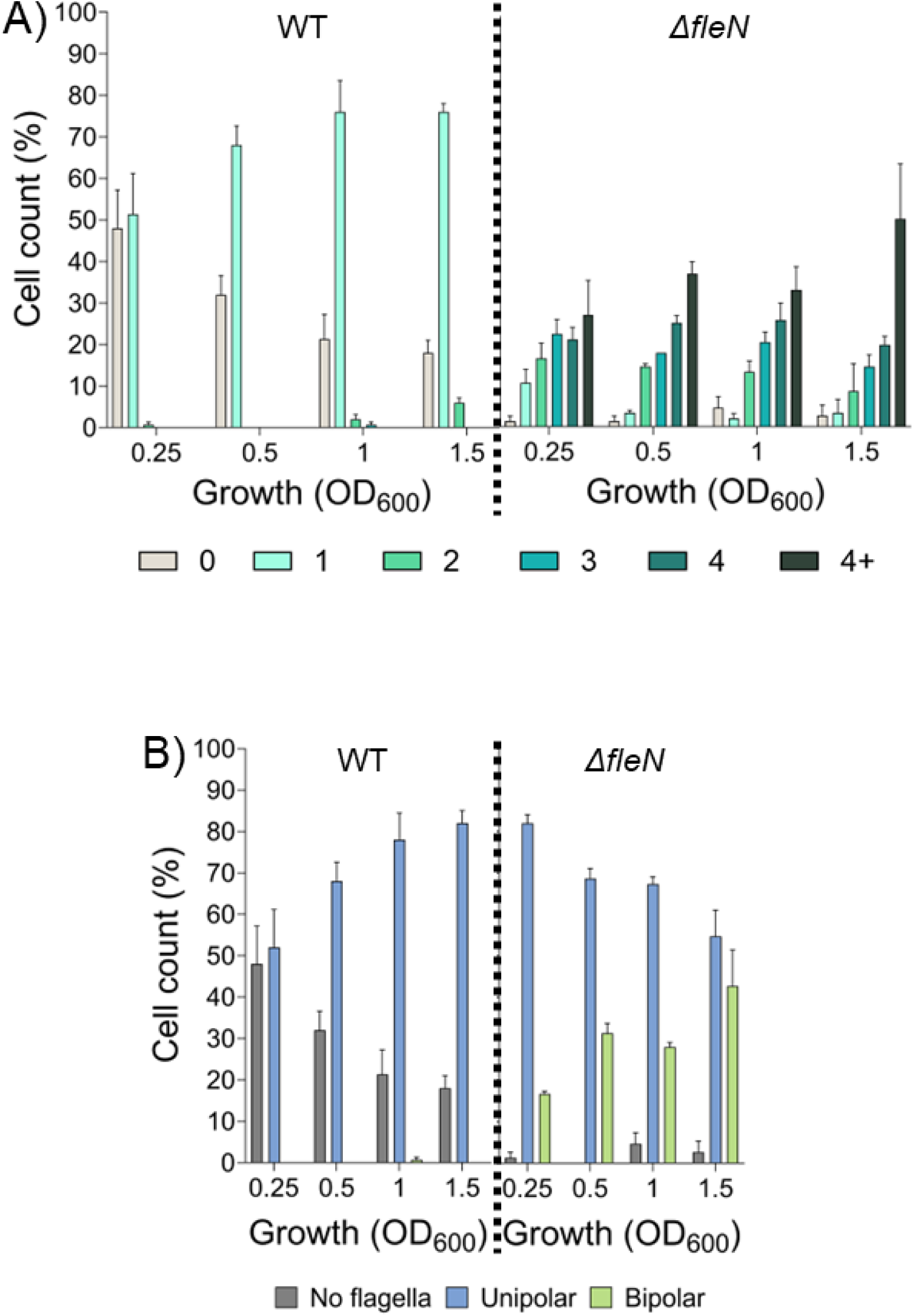
Monoflagellation is a robust trait in WT PA14 cells. A) Percentages of cells with different flagella numbers through early exponential (OD600 0.25), exponential (OD600 0.5), early stationary (OD600 1) and late stationary (OD600 1.5) of WT (left) and Δ*fleN* (left). Grey: aflagellated, Cyan: 1, Turquoise: 2, Dark turquoise: 3, Emerald green: 4, Dark green: More than 4 flagella per cell, 50 cells per replica, n=3. B) Percentage of cells with different flagellar placement from the cells in panel A), WT (left) and Δ*fleN* (left).

**Supplemental Fig.2:**
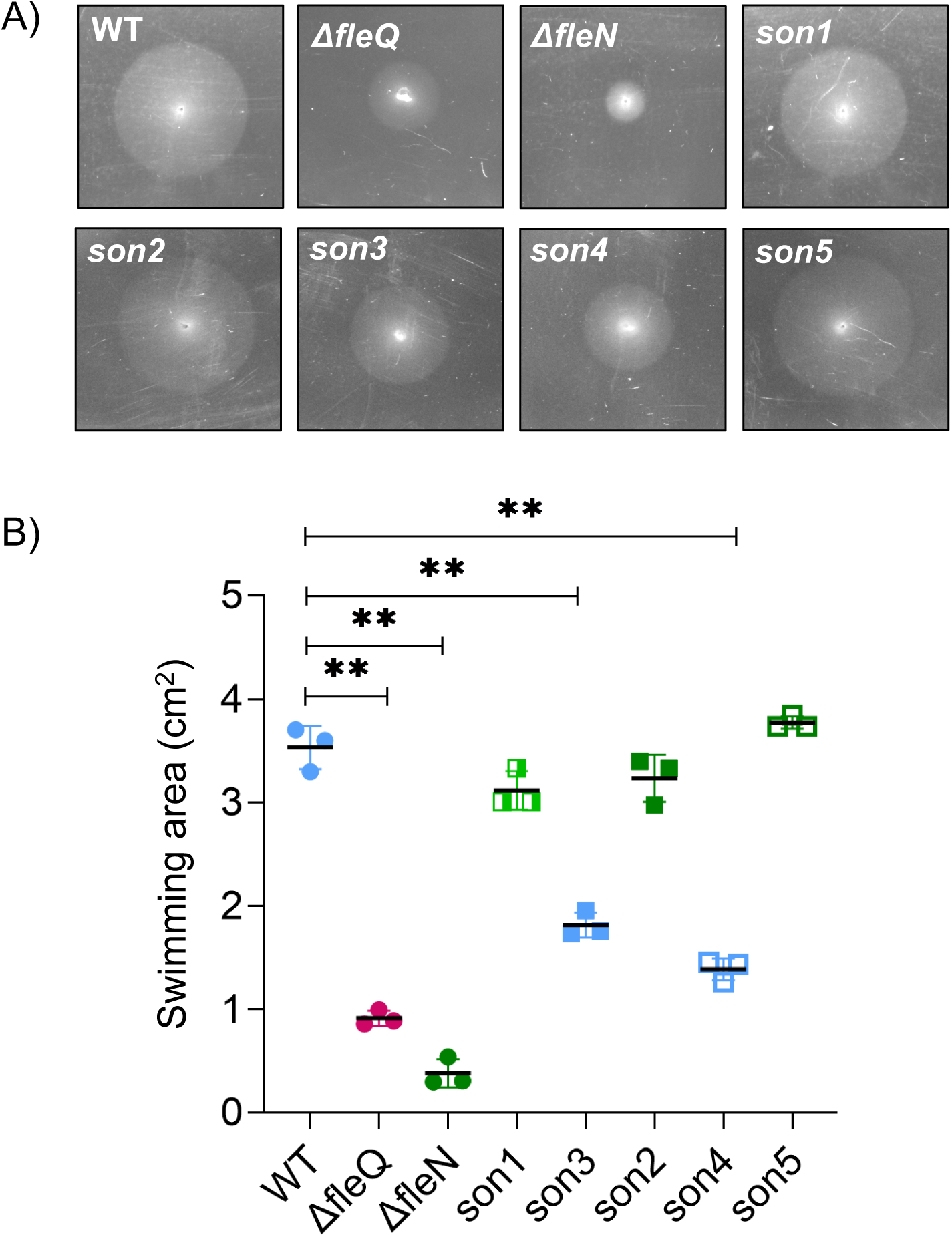
Swimming area of fleQ mutants. Swimming assay of WT, Δ*fleQ*, Δ*fleN* and WT background cells carrying the FleQ point mutations from the Δ*fleN* suppressors. n=3. B) Measure area of swimming from A).

**Supplemental Fig.3:**
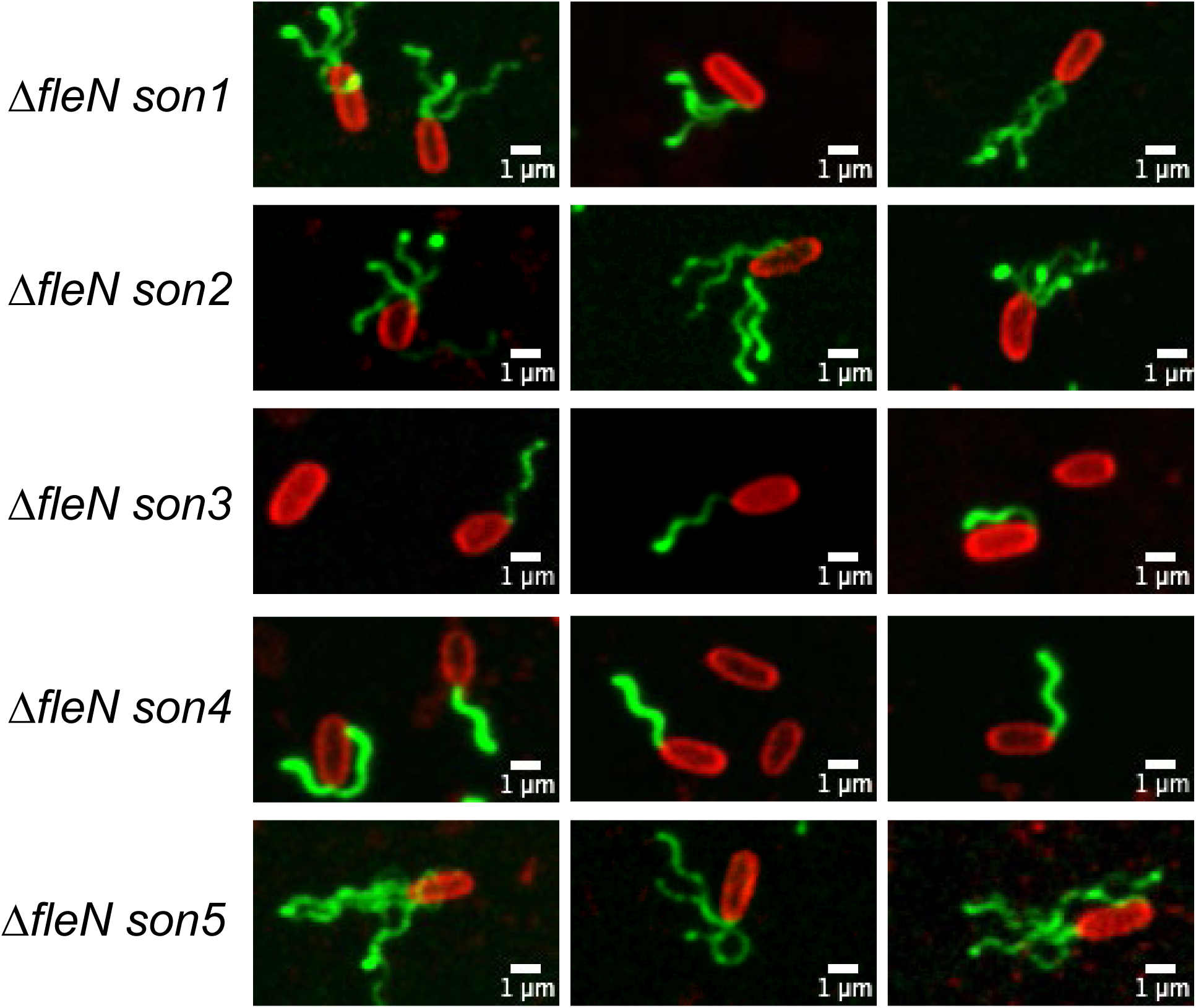
Flagella (Alexa488) and membrane staining (FM464) of suppressors of Δ*fleN*.

**Supplemental Fig.4:**
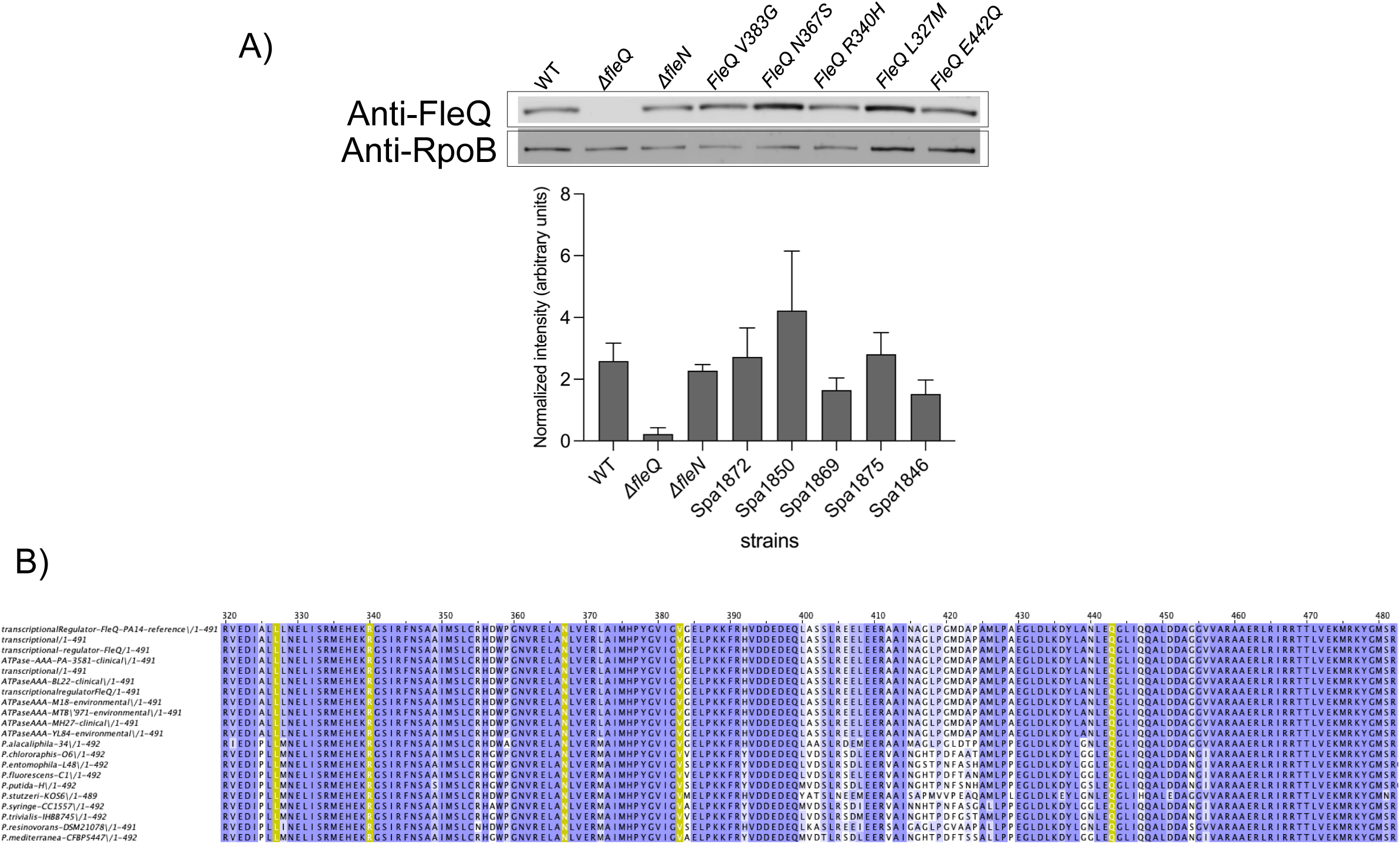
FleQ point mutations does not compromise stability of the protein. A) Representative image of Western Blot of *fleQ* mutants (upper panel) and its quantification using ImageLab (lower panel), n=3. B) Alignment of *fleQ* AAA+-HTH domains amino acid sequence from different types of Pseudomonades (single and multiflagellated strains). In yellow, the *fleQ* mutations found in the suppressors of Δ*fleN* in PA14.

**Supplemental Fig.5:**
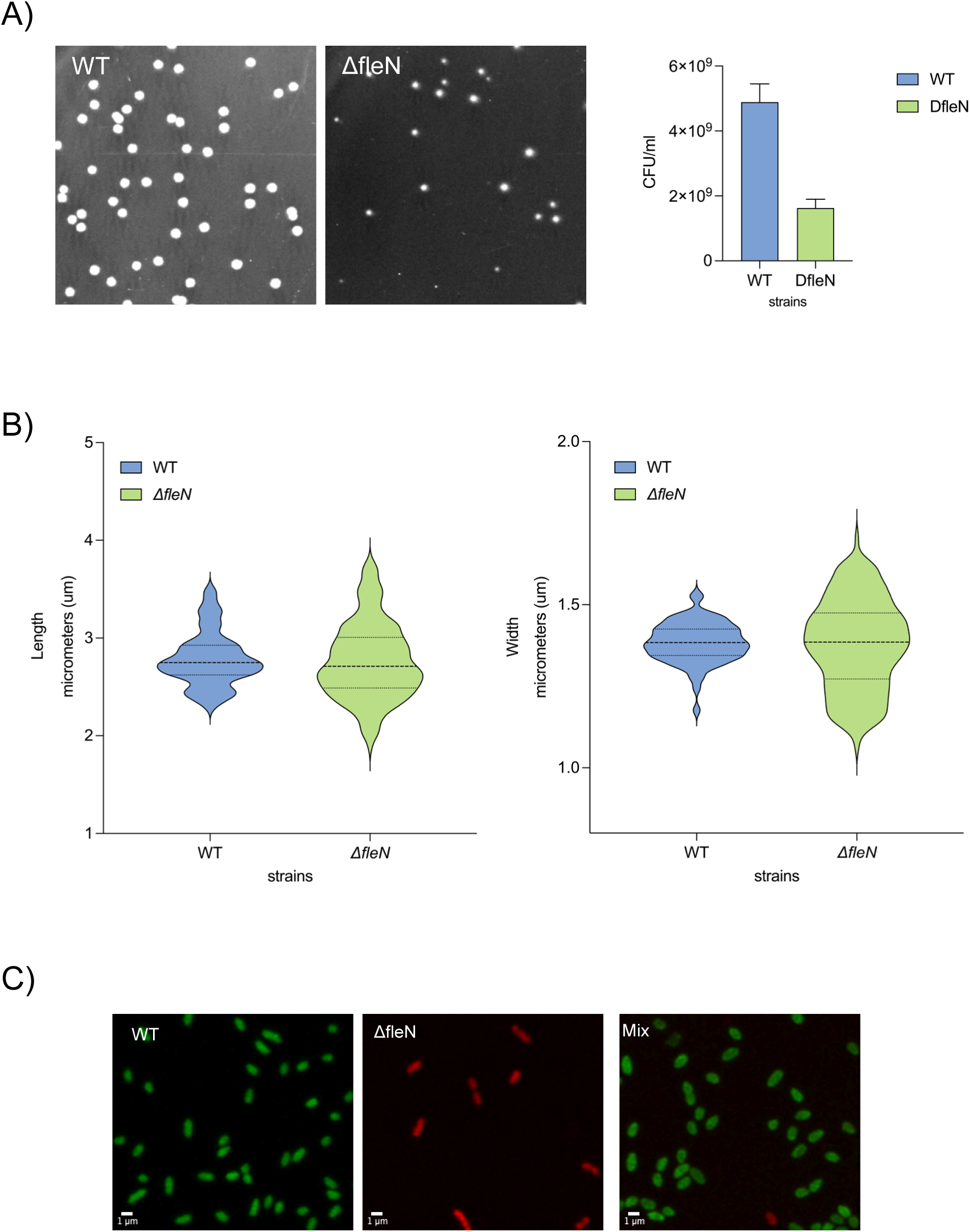
A) Independent growth of WT and Δ*fleN* cells for 20hrs at 37C in liquid, starting at the same OD600 (0.5). Colorimetric image of recovered colonies of each strain after 20hrs (Left) and its respective quantification of CFU/ml (right), n=3. B) Length (left) and width (right) of WT and Δ*fleN* cells at OD600 0.5, 50 cells per replica, 3 replicas. C) Nutrient competition between WT (green) and Δ*fleN* (red) cells. Representative image at the end of the competition (after 20hrs).

**Supplemental Fig.6:**
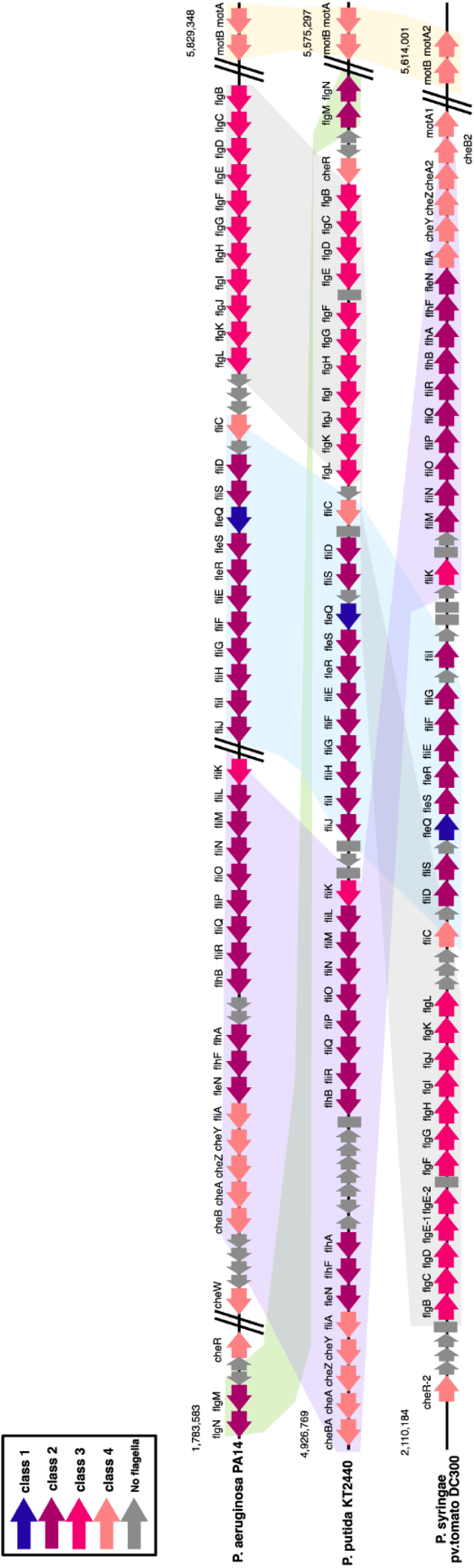
Synteny of flagellar genes between *P. aeruginosa* PA14, *P. putida* KT2440 and *P. syringae pv. tomato* DC300. Hierarchical flagellar classes are denotated by the color of the arrows.

**Supplemental Fig.7:**
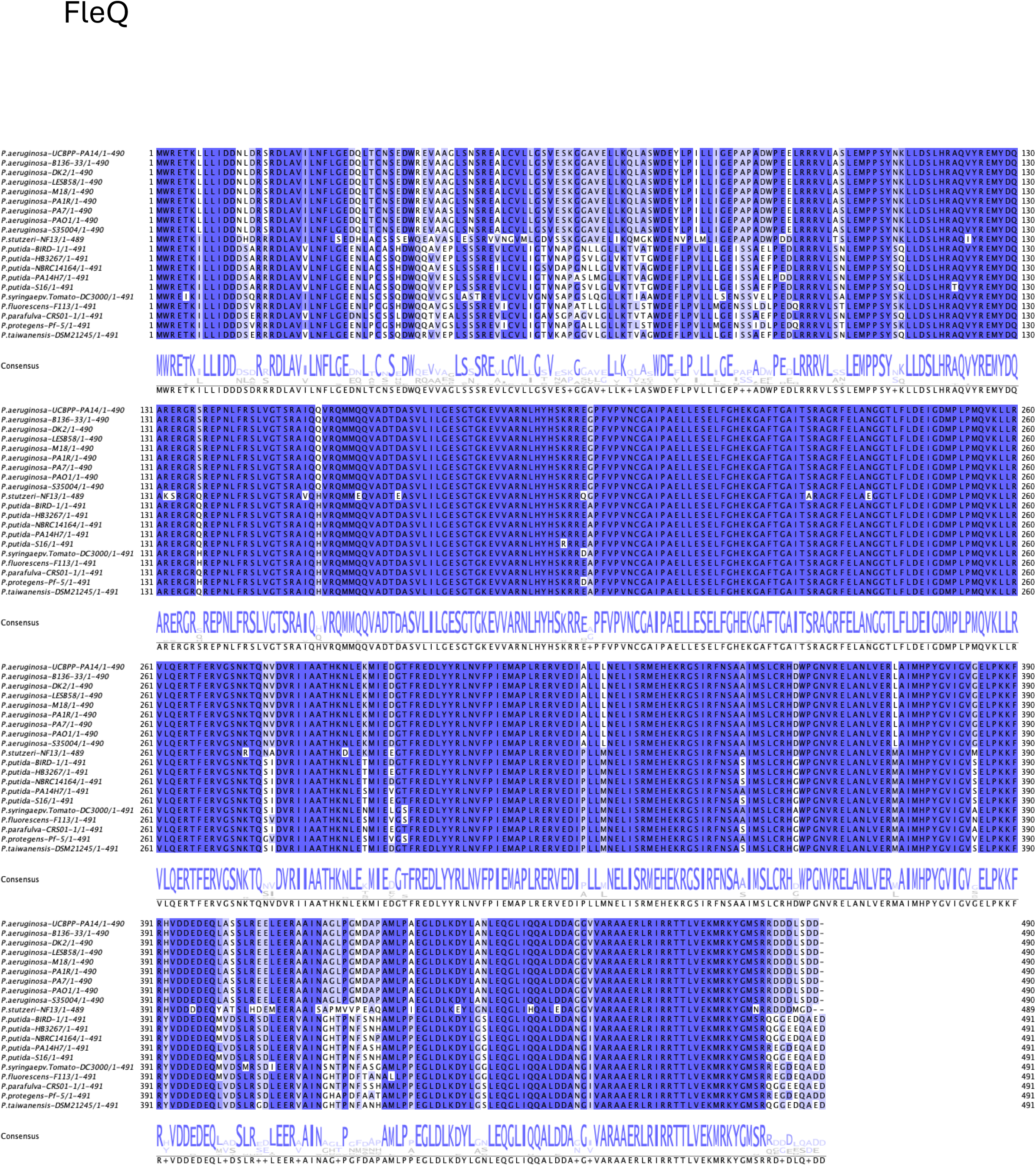
Alignment of FleQ amino acid sequence from different types of Pseudomonades (single and multiflagellated strains).

**Supplemental Fig.8:**
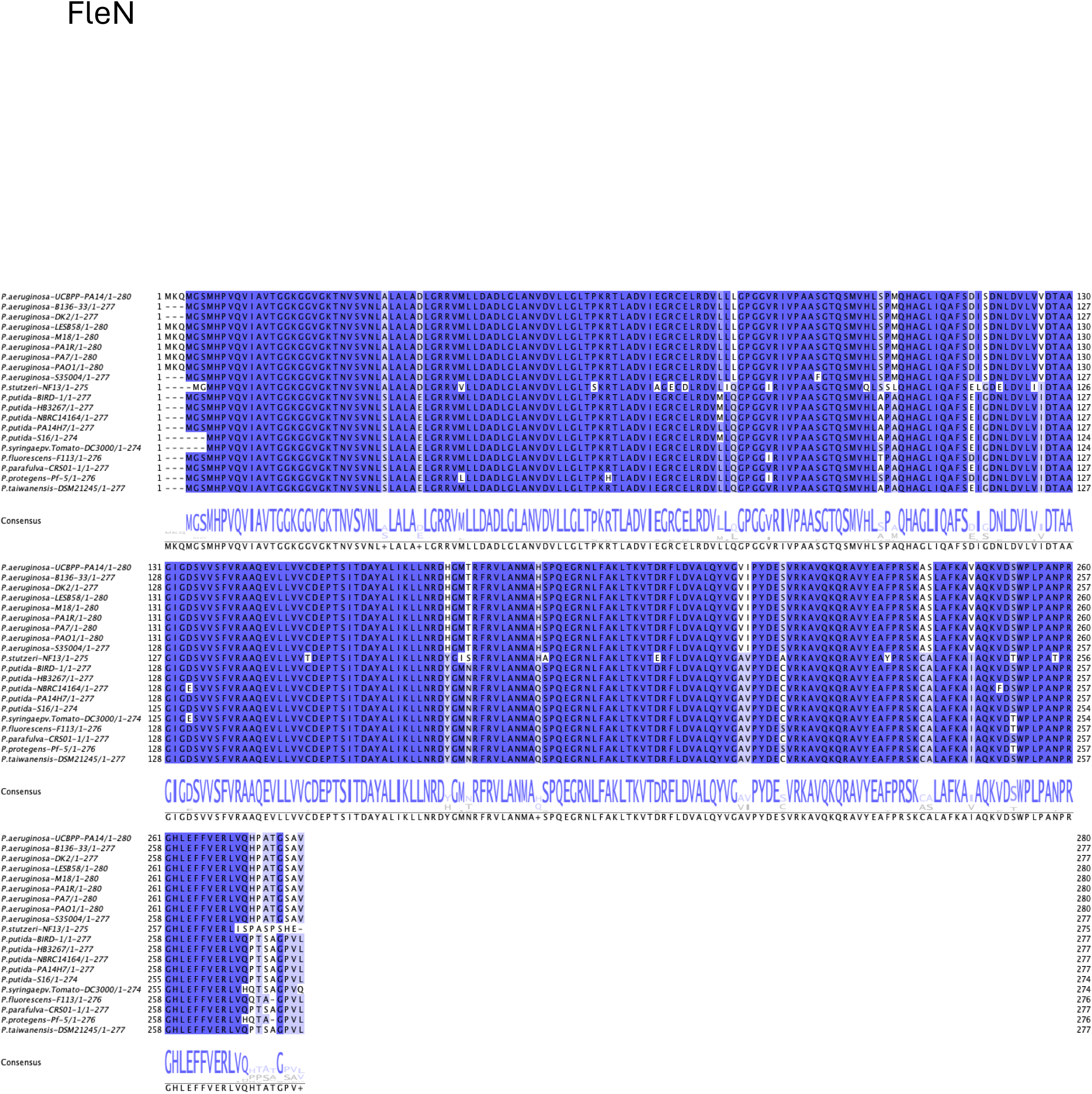
Alignment of FleN amino acid sequence from different types of Pseudomonads (single and multiflagellated strains).

**Supplemental Fig.9:**
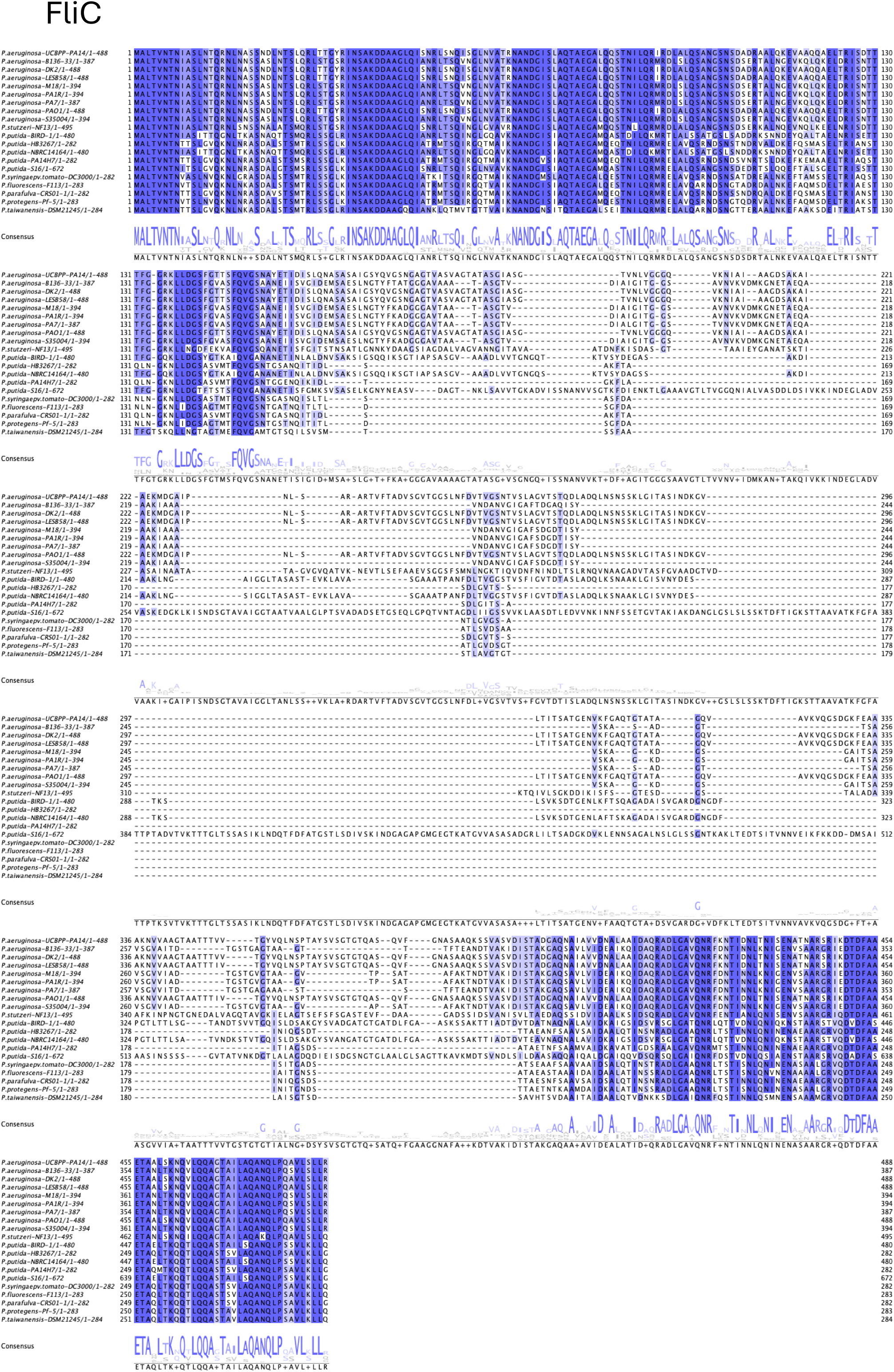
Alignment of FliC amino acid sequence from different types of Pseudomonads (single and multiflagellated strains).

